# Geometric Multidimensional Representation of Omic Signatures

**DOI:** 10.64898/2026.01.26.701791

**Authors:** Higor Almeida Cordeiro Nogueira, Enrique Medina-Acosta

**Affiliations:** Laboratório de Biotecnologia, Centro de Biociências e Biotecnologia, Universidade Estadual do Norte Fluminense, Brazil

**Author notes:** **Corresponding author**: Enrique Medina-Acosta, Address: Laboratório de Biotecnologia, Centro de Biociências e Biotecnologia, Universidade Estadual do Norte Fluminense, Avenida Alberto Lamego 2000, Parque Califórnia, Campos dos Goytacazes, RJ, CEP 28015-602, Brazil. Authors’.

**Keywords:** geometric representation, latent space, metabolic regulation, multi-omics, regulatory circuitries

## Abstract

Multi-omic signatures are widely used in biomarker discovery, precision oncology, and systems biology, yet they are typically treated as vectors or composite scores that collapse intrinsically multidimensional biological organization into one-dimensional summaries. As a result, their internal structure, contextual dependencies, and mechanistic coherence remain largely inaccessible. Here, we introduce a geometric framework that reconceptualizes omic signatures as multidimensional informational entities whose biological meaning arises from structural organization rather than molecular membership alone. Each signature is embedded in a shared latent space integrating regulatory, phenotypic, microenvironmental, immune, and clinical constraints, and represented as a convex polytope. This representation preserves internal organization and enables intrinsic geometric measurements—including barycenter distance, volume, anisotropy, and asymmetry—that quantify concordance, divergence, and latent complexity. We apply this framework to 24,796 metabolic regulatory circuitries reconstructed across 32 TCGA cancer types, encoded as paired regulatory and metabolic signatures in an 18-dimensional latent space. Geometric analysis shows that discordance predominates: most circuitries occupy strong or extreme discordance regimes and display high-dimensional, frequently asymmetric geometries, whereas fully concordant circuitries are rare and structurally constrained. These geometric phenotypes stratify metabolic pathways and superfamilies in reproducible, non-uniform patterns that are not detectable with vector- or network-based representations. By transforming omic signatures into measurable geometric objects, this framework enables principled comparison, de-redundancy, and mechanistic interpretation of multi-omic biomarkers, providing a scalable approach for analyzing complex regulatory systems across cancer and beyond. All geometric representations and derived descriptors are available through the SigPolytope Shiny application (https://sigpolytope.shinyapps.io/geometricatlas/).

## 1. Introduction

### 1.1. Limits of Vectorial Representations in Multi-Omic Biomarker Research

The prevailing paradigm in candidate biomarker cancer research treats omic associations as fundamentally vectorial relationships, in which a molecular measurement at a single omic layer is statistically linked to a clinical or phenotypic attribute (1–3). While vectorial representations are indispensable as computational tools, their use as final biological abstractions collapses the biological complexity of tumor systems into one-dimensional descriptors that cannot capture mechanistic organization, conditionality, or multi-layer coherence. Even when the field adopts the language of “signatures,” such entities are typically handled as simple lists, scores, or co-expressed gene vectors rather than as structured informational objects with internal organization or topology (1, 4–6).

In practice, however, omic signatures are clinically motivated constructs designed to stratify patients, predict outcomes, assess risk, or guide therapeutic decision-making. Their biological and clinical meaning, therefore, arises not from isolated measurements but from the integration of multiple informational dimensions (7–10). Viewed in this way, signatures function as biomarker objects that organize complex multi-omic associations into a form capable of supporting prediction, stratification, and phenotypic interpretation.

To address this complexity, we previously introduced a more expressive nomenclature to encode multi-omic signatures by explicitly mapping their constituent associations across biological layers, mechanisms, and phenotypes (11) (Figure 1A). These symbolic representations provide a useful linguistic scaffold, allowing readers to trace which omic layers connect to which phenotypic axes, and they support structured interpretation through dedicated tools such as the CancerRCDShiny browser (https://cancerrcdshiny.shinyapps.io/cancerrcdshiny/) and the Multi-omic OncometabolismGPS Shiny (https://oncometabolismgps.shinyapps.io/Multi-omicOncometabolismGPSShiny/). Importantly, this nomenclature also enables the formal pairing of signatures into regulatory circuitries, illustrated schematically in Figure 1B.

**Figure 1.**
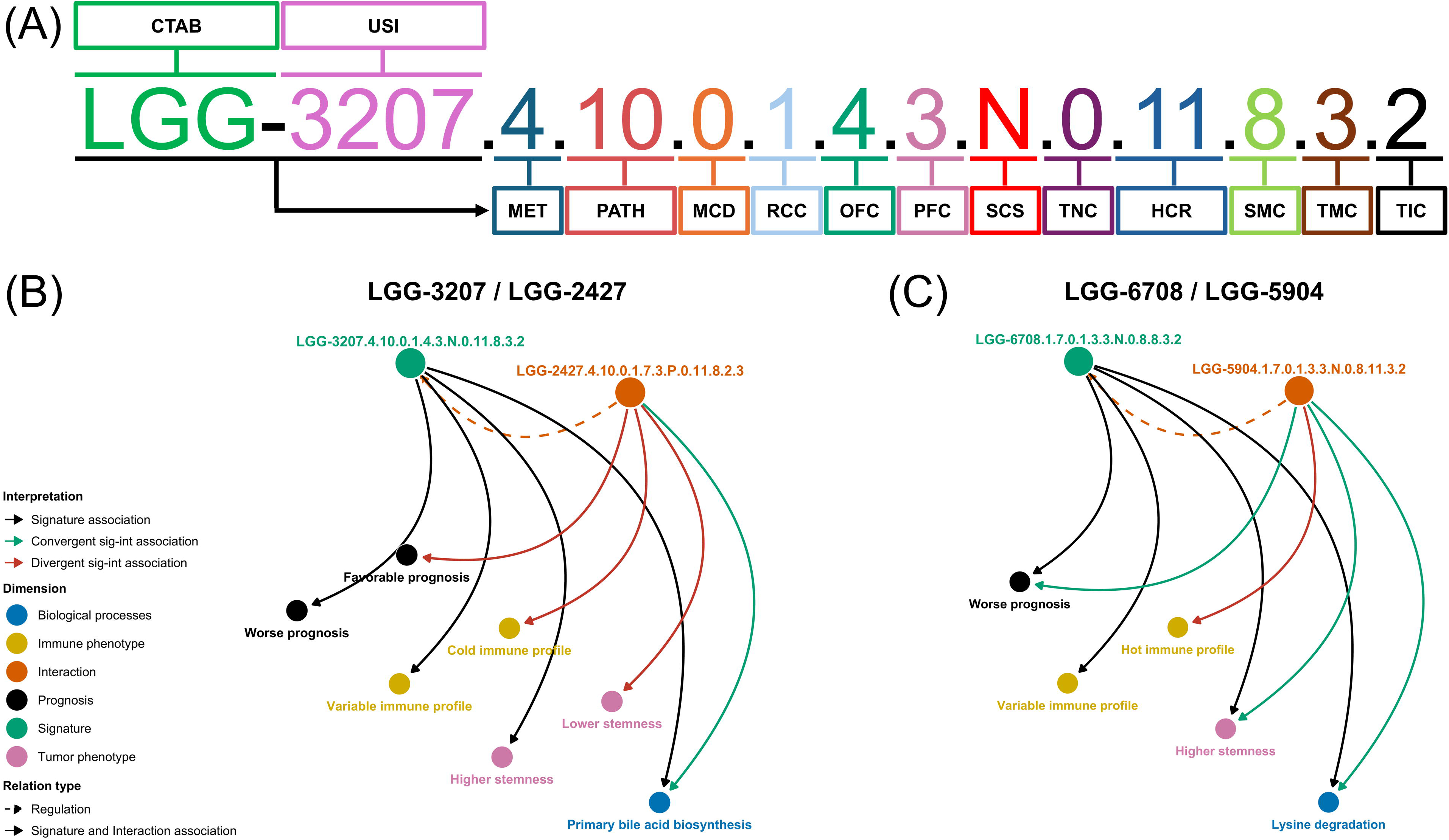
Nomenclature of multi-omic metabolic signatures and examples of regulatory circuitry configurations. (A) Symbolic encoding of multi-omic metabolic signatures. Panel A summarizes the nomenclature system previously introduced to encode multi-omic metabolic signatures in cancer(11). Each signature is represented by an ordered, multi-token alphanumeric identifier that integrates tumor context, metabolic class and pathway assignment, regulatory associations, omic and phenotypic layers, expression behavior, survival impact, tumor–microenvironmental context, and immune state. This encoding provides a structured annotation scheme for the regulatory circuitry datasets generated in prior work. A concise description of the nomenclature fields and encoding logic is provided in the Supplementary Material. (B) Divergent metabolic regulatory circuitry. Panel B illustrates a representative divergent regulatory circuitry drawn from the previously generated circuitry dataset and used here for geometric analysis. In this configuration, the paired metabolic and regulatory signatures exhibit opposing directions across one or more biological dimensions—such as phenotypic associations, survival risk profiles, or immune context—despite being linked within the same tumor type. This divergence reflects regulatory–metabolic misalignment rather than coordinated action. In the present study, such divergent circuitries are embedded into a shared latent space to quantify geometric separation and directional discordance between their component signatures. (C) Convergent metabolic regulatory circuitry. Panel C illustrates a representative convergent regulatory circuitry in which the paired metabolic and regulatory signatures align consistently across molecular, phenotypic, clinical, and immune dimensions. In this configuration, both components encode concordant directionality with respect to tumor–normal behavior, survival associations, and microenvironmental or immune context, reflecting coordinated regulatory control of the metabolic program within the tumor. Convergent circuitries represent cases of regulatory agreement rather than opposition and provide a conceptual contrast to divergent configurations, serving as a reference point for interpreting geometric concordance in the latent space.

Nevertheless, such nomenclatures and interpreter tools remain fundamentally relational. They function as decoders that convey which components connect, but not how strongly, in what direction, or with what emergent organization. As a result, they preserve semantic meaning and improve accessibility but do not retain heterogeneity, scale, anisotropy, or geometric structure. These approaches, therefore, remain inherently lossy representations—valuable for navigation and interpretation, yet insufficient to preserve the full multidimensional organization of omic signatures. For example, two signatures with opposite survival directionality or tumor–normal behavior may appear highly similar when reduced to gene overlap, list-based annotations, or composite scores, despite encoding fundamentally divergent biological and clinical implications. This limitation motivates the need to move beyond symbolic or interaction-based encodings toward representations that honor the intrinsic multidimensional nature of omic signatures as informational entities.

The geometric framework introduced here is applied, at scale, to a large collection of metabolic regulatory circuitries reconstructed across multiple cancer types. These circuitries, together with their biological annotation and population-level organization, are described in a companion atlas manuscript (11). In the present work, this atlas serves as an empirical substrate for evaluating the representational and analytical properties of the geometric formalism under realistic, heterogeneous, multi-omic conditions. Importantly, this manuscript is not intended as an extension of that biological atlas, nor as a catalog of novel regulatory interactions. Rather, it establishes a generalizable representational framework for omic signatures, independent of any specific pathway, disease context, or biological domain, using atlas-scale data solely to demonstrate how geometric representation enables principled comparison, de-redundancy, and mechanistic interpretation beyond conventional vector- or network-based approaches.

Importantly, the present study is not intended as a biological atlas extension or as a catalog of newly discovered regulatory interactions. Rather, it is a conceptual and methodological contribution that addresses a more fundamental problem in contemporary bioinformatics: how intrinsically multidimensional omic signatures should be represented, compared, and interpreted once they move beyond simple vectors or lists. We use a large collection of previously reconstructed metabolic regulatory circuitries as an empirical substrate to stress-test this representational problem under realistic, heterogeneous, multi-omic conditions. In this sense, the atlas-scale application serves as a proving ground rather than as the primary objective. The central aim of this work is to establish a generalizable geometric formalism for omic signatures, independent of any specific pathway, disease context, or biological domain, and to demonstrate how such a formalism enables forms of comparison, de-redundancy, and mechanistic reasoning that are inaccessible to conventional vector- or network-based representations.

### 1.2. Signatures: From Operational Descriptors to Structured Biological Entities

Although the term signature is now ubiquitous in biomarker discovery, its usage in the literature is often imprecise, frequently referring to any statistical association between a set of molecular features and a phenotype. Importantly, this critique does not question the clinical utility of established signatures, but rather the conceptual framework used to interpret their biological meaning. In this reductionist framing, signatures function primarily as correlation-derived lists or composite scores, lacking explicit mechanistic architecture, contextual grounding, or internal informational structure. Operationally, a gene signature is typically defined as a gene or specific set of genes whose expression patterns collectively characterize a biological state, cancer subtype, prognosis, or treatment response. For example, the PAM50 classifier, a 50-gene signature, stratifies breast cancers into Luminal A, Luminal B, HER2-enriched, and Basal-like intrinsic subtypes (12). Similarly, the Oncotype DX 21-gene signature predicts recurrence risk and chemotherapy benefit in tamoxifen-treated, node-negative breast cancer (13), while the T cell–inflamed gene expression profile (GEP), composed of IFN-γ–responsive genes, predicts response to PD-1 blockade (14).

While these signatures are intended to function as biological barcodes, their conceptual definition often remains loose. Most reported signatures aggregate molecular signals without explicitly encoding coordinated biological behavior, and may or may not correspond to coherent regulatory programs. In principle, a signature should serve as a distinct and specific, even unique, molecular fingerprint; in practice, however, it rarely meets this criterion. Consequently, signatures are frequently interpreted as mechanistically meaningful entities when they remain, at their core, operational descriptors with limited internal structure, lacking explicit representation of coordinated biological organization (15–17).

Recognizing this discrepancy is critical. Without a rigorous definition that frames signatures as multidimensional informational constructs—rather than residual statistical associations—the field risks overestimating their mechanistic significance while underestimating the conceptual and analytical rigor required for developing clinically credible biomarkers.

An omic signature, however, is neither a gene list, nor a panel, nor a simple vector. By its nature, it is a multidimensional informational construct that emerges from coordinated molecular relationships spanning regulatory hierarchies and contextual phenotypes. We therefore define an omic signature as a structured informational entity composed of an integrated set of biomolecular components—including genes, transcripts, proteins, regulatory RNAs, and epigenomic or genomic alterations—whose organization transcends mere co-expression. Rather than reflecting isolated correlations, an omic signature arises from the systematic integration of biological layers, mechanistic axes, and phenotypic contexts.

Each signature thus synthesizes consistent relationships across omic dimensions, biological mechanisms, and tumor phenotypes, forming a structured evidence space that captures the functional state of cellular processes within a defined tissue, cancer type, or clinical condition. Whereas unitary biomarkers encode linear, unidimensional associations, an omic signature represents a reproducible, high-dimensional informational field, enhancing predictive performance, mechanistic interpretability, biological stratification, and translational applicability. These considerations motivate a deeper conceptual reformulation of what an omic signature fundamentally is and how its informational structure should be represented.

### 1.3. Redefining Omic Signatures as Emergent Multi-Informational Entities

Building upon our prior work(11), we focus here on metabolic regulatory circuitries, a biologically grounded construct defined as a paired entity composed of two interacting signatures: a metabolic signature capturing the functional metabolic program of a tumor context, and a regulatory signature capturing the upstream molecular influences that modulate that program (Figure 1B). This paired formulation was introduced to encode regulatory alignment and opposition across molecular, phenotypic, immune, and clinical dimensions, and has been systematically reconstructed at atlas scale. Importantly, the present study does not seek to re-establish the biological properties of these circuitries, which are reported elsewhere(11). Instead, we use them as an empirical proving ground to address a distinct and unresolved question: how intrinsically multidimensional, paired entities should be represented, compared, and interpreted. Because regulatory circuitries are defined by internal structure, directionality, and potential discordance between their constituent signatures, they provide a uniquely suitable substrate for developing and validating a framework that preserves multidimensional organization rather than collapsing it into vectorial summaries.

A rigorous redefinition of omic signatures, therefore, requires moving beyond the classical paradigm that represents a signature as a solitary vector of measurements. This vectorial framing presumes linearity, independence among dimensions, and static interpretability— assumptions that rarely capture biological reality. In contrast, an omic signature should be understood as an n-dimensional, multi-informational entity, whose structure and function emerge from coordinated interactions across molecular, regulatory, and phenotypic layers. This reconceptualization aligns more faithfully with the heterogeneous, dynamic, and context-dependent nature of tumor systems, providing a more appropriate framework for representing the informational richness inherent in multi-omic data.

An omic signature integrates multiple informational modalities that differ in scale, generative mechanisms, and interpretive context. These include discrete alterations such as mutations and copy-number changes; continuous regulatory measurements such as mRNA, protein, and metabolite abundances; categorical state descriptors including stress responses, pathway activation states, and immune phenotypes; and topological attributes reflecting pathway membership, regulatory network connectivity, or interaction motifs. Each component contributes not a single scalar measurement but a distinct informational domain, characterized by its own scale, noise structure, and regulatory determinants. The identity of the signature therefore emerges from the coordinated interactions among these domains, collectively defining a coherent, multidimensional representation of biological state.

The defining property of an omic signature is thus its emergence: its biological meaning arises not from any individual feature, but from the organized behavior of all components operating under specific regulatory constraints. Rather than reflecting isolated correlations, a signature derives its identity from coordinated relationships across molecular, regulatory, and phenotypic dimensions, such that its meaning cannot be reduced to marginal measurements alone. Biologically, these relationships arise from diverse cross-layer interactions, including metabolite-driven rewiring of transcriptional programs, microRNA-mediated buffering of metabolic enzymes, context-specific immune modulation of metabolic demand, and stress-induced compensatory pathways characteristic of survival or regulated cell-death processes. In this view, a signature is defined by its internal organization and dynamic relationships, not merely by molecular membership.

This organizational structure fundamentally distinguishes omic signatures from traditional vectorial representations. Signatures that appear similar when reduced to gene lists or composite scores may encode profoundly different regulatory architectures, reflecting distinct patterns of inter-layer coupling, contextual constraint, or phenotypic engagement. Conversely, signatures with limited molecular overlap may converge toward similar biological behaviors when their internal organization is considered. Importantly, omic signatures are inherently contextual: their structure and interpretation are shaped by tissue identity, tumor lineage, microenvironmental pressures, therapeutic exposure, and perturbation history. As a result, the same nominal signature may exhibit distinct organizational behavior across biological settings, explaining the frequent lack of portability observed when signatures are treated as fixed vectors.

This reconceptualization carries important methodological implications. Predictive models must account for interaction structure rather than rely solely on marginal effects; evaluation frameworks must assess behavioral organization rather than single performance metrics; and interpretation must focus on latent structure and cross-layer dependencies rather than differential abundance alone. Within this paradigm, an omic signature is best understood as a structured, multidimensional informational entity whose biological meaning emerges from higher-order organization.

Building on this perspective, the multidimensional organization of omic signatures naturally motivates the use of geometric representations that preserve structure rather than collapse it. By representing signatures as geometric objects—specifically as convex polytopes rather than vectors—the field acquires both a language and a framework that more accurately reflect biological organization. Two signatures that appear similar at the gene-list level may yield markedly different geometric structures, revealing divergent mechanistic architectures or phenotypic trajectories, while signatures with distinct molecular compositions may converge toward similar geometries, suggesting functional redundancy or systems-level equivalence. The formal geometric framework underlying this representation is described in the Supporting Information.

### 1.4. Geometry as an Interpretive Lens for Omic Signatures

Once an omic signature is conceived as a multidimensional informational entity, geometric representation becomes a natural interpretive lens, as it preserves metric structure and enables quantitative comparison of organization rather than connectivity alone. When signatures are viewed only as lists or vectors, their internal organization remains opaque; geometric representation makes this organization directly visible by transforming relationships into spatial structure. In this framework, the three-dimensional convex hull serves as an accessible conceptual projection, revealing latent structural relationships—emerging from integrated regulatory, phenotypic, and mechanistic constraints encoded by the latent axes—that encode coordination, divergence, and fragmentation into subprograms among molecular components, rather than imposing an artificial geometry. The hull delineates the minimal envelope defined by the spatial organization of components within latent space, allowing multidimensional structure to be apprehended visually and conceptually (schematically illustrated in Figure 2A; an interactive HTML version used during analysis and exploratory inspection is provided in the Supporting Information, Figure 2_HTML). Figure 2 is intended exclusively as a schematic aid and does not depict any empirical circuitry or latent embedding used in subsequent analyses. Panels B–F then illustrate idealized canonical organizational regimes, which later recur in the analysis of real omic signatures, ranging from coherent single-axis structure to heterogeneous modular organization.

**Figure 2.**
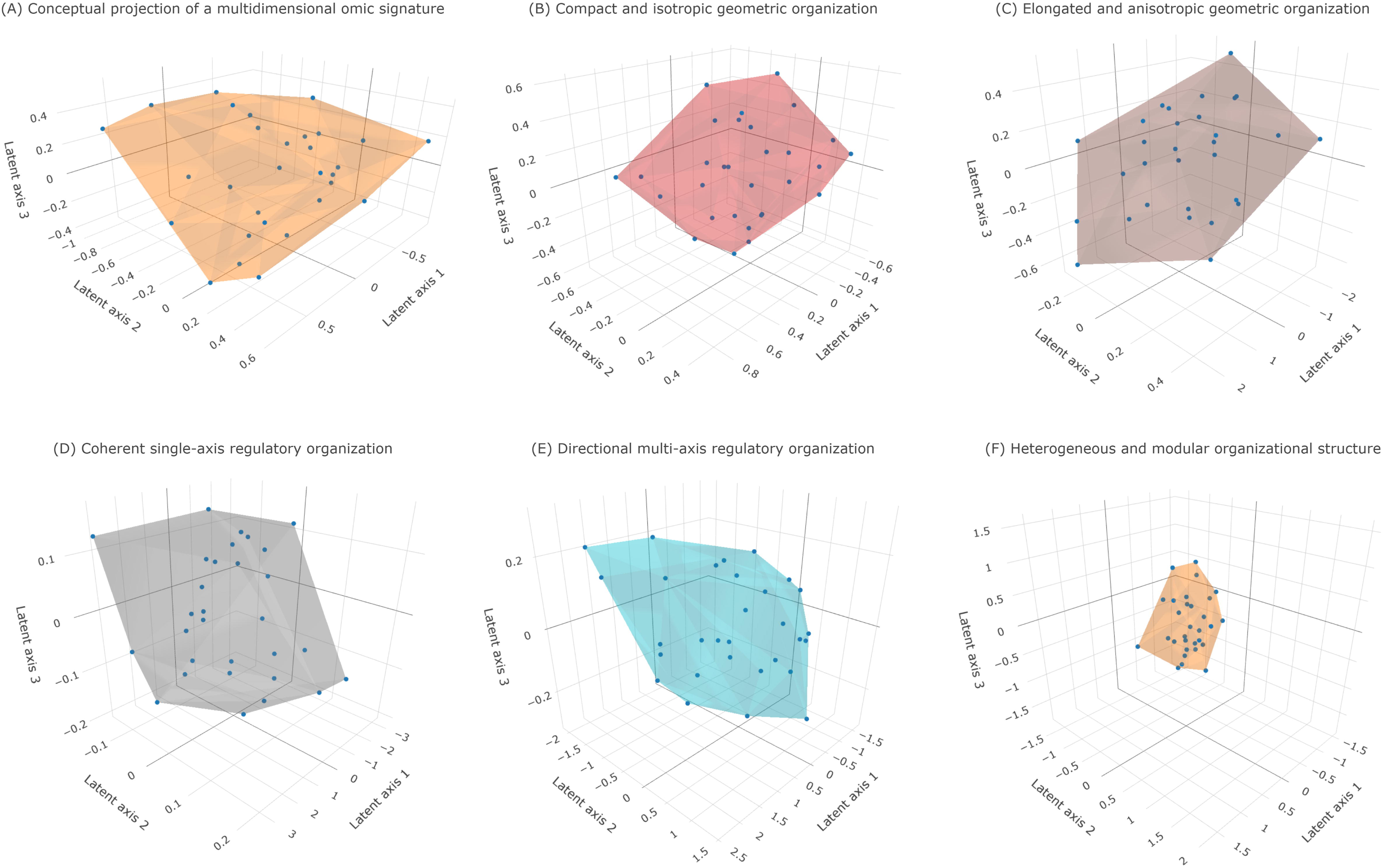
Conceptual geometric representations of omic signature organization based on simulated data. (A) Schematic illustration of an idealized omic signature represented as a multidimensional informational entity projected into a three-dimensional latent space. All points shown correspond to synthetically generated components (n = 30 points; convex-hull vertices = 16) and do not represent empirical molecular measurements. Points are positioned to illustrate hypothetical contributions across mechanistic, phenotypic, and contextual dimensions. The convex hull delineates the minimal geometric envelope enclosing the component cloud, providing a qualitative visualization of higher-dimensional organizational structure. (B–C) Conceptual examples of distinct geometric fingerprints arising from different internal organization in simulated signatures. Although the signatures contain comparable numbers of components (n = 30 points each), differences in spatial arrangement yield compact, isotropic geometries (B; hull vertices = 16) or elongated, directional geometries (C; hull vertices = 15), illustrating coherent versus axis-dominated organizational patterns, respectively. (D–F) Canonical geometric regimes illustrating organizational heterogeneity using synthetic point clouds. Compact and isotropic hulls (D; hull vertices = 13) indicate coherent, single-axis– dominated behavior; elongated and anisotropic hulls (E; hull vertices = 17) reflect directional structure and partial divergence; irregular and faceted hulls (F; hull vertices = 14) indicate heterogeneous organization consistent with the coexistence of multiple semi-independent biological modules. All geometries shown are generated exclusively from simulated data and are intended solely to illustrate conceptual principles of geometric organization in multidimensional omic signature space. Axis orientations, relative scales, and tick values serve as illustrative cues only and do not correspond to empirical measurements, enforced metric distances, or real omic coordinate systems. These representations do not depict experimental results but provide schematic guidance for interpreting geometric patterns in subsequent analyses of real data. (Source code and interactive HTML versions are provided in the Supporting Information.)

Within this representation, each molecular component corresponds to a point whose position reflects its contribution across mechanistic, phenotypic, and contextual dimensions (Figure 2A). The collective distribution of these points forms a molecular cloud, and the enclosing convex hull constitutes a geometric fingerprint of the signature. This fingerprint is intrinsic: signatures with distinct internal organization yield distinct geometric forms, even when they share comparable numbers of components or exhibit overlapping phenotypic associations, as illustrated by compact, isotropic geometries versus elongated, anisotropic geometries (Figure 2B–C).

The geometry of this envelope carries direct biological meaning. Compact and isotropic forms indicate coherent organization, consistent with convergence toward a dominant regulatory or phenotypic axis (Figure 2B; Figure 2D). In contrast, elongated or anisotropic geometries reveal directional structure, reflecting axis-dominated organization, partial divergence, or hierarchical regulation across latent dimensions (Figure 2C; Figure 2E). More irregular and faceted geometries expose heterogeneous internal organization, implying the integration of multiple, potentially semi-independent biological modules within a single signature (Figure 2F). In this sense, geometric form functions as a visual and conceptual map of biological organization, where distance encodes discordance, volume reflects latent complexity, and shape captures organizational coherence.

Importantly, this interpretive capacity does not depend on specific molecular identities but on structural relationships among components. Geometry, therefore, enables signatures to be compared, interpreted, and contextualized based on organization rather than composition alone. By revealing coherence, directionality, and heterogeneity directly from structure, geometric representation provides an interpretive layer that complements—but cannot be replaced by—vector-based summaries or list-based annotations.

## 2. Materials and Methods

### 2.1. Conceptual Framework for Geometric Representation of Omic Signatures

Building upon the geometric rationale introduced above, we represent each omic signature as a finite set of biomolecular components embedded within a shared latent space, where each latent dimension encodes a biologically meaningful attribute, including regulatory effect, phenotypic breadth, contextual stability, or mechanistic contribution. In this framework, individual molecular components are mapped to latent coordinates that summarize their integrated multi-omic associations, enabling signatures to be treated as structured geometric objects rather than unordered molecular lists.

Within this latent space, the structural boundaries of a signature are defined by its convex envelope, corresponding to the minimal region enclosing all constituent components while preserving their multidimensional organization. This envelope provides a topology-preserving representation of internal structure and enables direct geometric interrogation of signature organization. For visualization and comparative analyses, the latent representation is projected into three dimensions while retaining the salient geometric relationships that characterize each signature. Importantly, barycenter position captures only the net directional tendency of a signature in latent space, whereas the convex hull encodes its internal dispersion, anisotropy, and multi-axis deformation—structural properties that are invisible to distance-based analyses, clustering, or PCA coordinates alone. Throughout the manuscript, we use convex polytope to refer to the intrinsic n-dimensional geometric object defined in latent space, and convex hull to denote its minimal enclosing envelope or its three-dimensional projection used for visualization and qualitative inspection.

From this representation, we derive a family of intrinsic geometric measurements that extend beyond conventional association statistics. The volume of the convex envelope serves as a proxy for multidimensional amplitude, reflecting the breadth of biological programs engaged by the signature. The orientation of principal axes captures dominant sources of mechanistic variation, while anisotropy distinguishes signatures driven by a single coherent process from those reflecting multiple, partially orthogonal subprograms. Additional descriptors—including centroid location, spatial extent, surface morphology, and angular divergence between signatures—enable systematic comparison, classification, and stratification.

Because these measurements arise from structural organization rather than molecular identity, the framework is inherently generalizable. It is independent of any specific pathway, phenotype, or disease model and is therefore applicable beyond cancer to any biological context in which omic signatures are used to encode mechanistic hypotheses, stratification schemes, or predictive biomarkers.

By treating signatures as geometric objects rather than vectorial summaries, this approach reframes biological inference as a problem of structural quantification. Signatures can be compared, ranked, merged, or excluded on principled geometric grounds, providing a robust basis for de-redundancy, mechanistic discrimination, and interpretability. In this formulation, geometry constitutes a methodological foundation rather than a visualization accessory, rendering multi-omic signatures analytically comparable across datasets, biological contexts, and disease areas. Formal mathematical definitions and computational implementations underlying this framework are provided in the Supporting Information.

### 2.2. Geometric Representation of Regulatory Circuitries

To capture the multidimensional behavior of regulatory circuitries beyond conventional one-dimensional summary statistics, we embedded each circuitry into a shared latent geometric space and derived polytope-based descriptors integrating correlation structure, tumor–normal shifts, survival associations, tumor microenvironment (TME), and immune contexture.

The regulatory circuitry atlas used in this study (11) exhibits broad Pan-Cancer coverage, encompassing circuitries derived from 32 distinct TCGA cancer types, including solid tumors and hematological malignancies. These cancer types span diverse tissue origins and biological contexts, ranging from central nervous system tumors and endocrine neoplasms to gastrointestinal, gynecological, urological, and immune-related cancers. This extensive coverage ensures that the atlas provides a comprehensive foundation for the geometric analyses performed here and is not dominated by a narrow subset of tumor contexts.

Each regulatory circuitry was represented as a paired entity composed of two latent profiles: one corresponding to the regulatory signature side (sig) and one to the interacting partner side (int). For each side, we constructed an 18-dimensional latent vector aggregating: (i) the Spearman rank correlation coefficient (ρ) between omic-layer–derived variables and their associated phenotypic attributes, together with the corresponding multiple-testing–adjusted P value; (ii) the direction of tumor-versus-normal differential expression and its −log10 Wilcoxon P value; (iii) Cox regression direction (risk versus protective) and associated P values for four survival endpoints (overall survival, disease-specific survival, disease-free interval, and progression-free interval), together with their corresponding log-rank χ² statistics; and (iv) a continuous microenvironment score and a categorical immune-class assignment. Categorical variables—including tumor–normal status, survival effect (risk, protective, or non-significant), and immune class (immune-hot, cold, excluded, or intermediate)—were consistently encoded as signed numerical values (−1, 0, +1). When a survival endpoint was non-significant, the associated direction, strength, and χ² contributions were explicitly set to zero to prevent non-informative axes from deforming the circuitry geometry.

Latent vectors from all circuitries (2N total vectors: N circuitries × 2 sides) were jointly embedded using principal component analysis (PCA). Before embedding, each latent dimension was globally centered and scaled across all circuitries.

In this context, the principal axes of the geometric space are defined operationally as the leading principal components of the latent tensor. Latent axes summarize integrated variance patterns emerging from regulatory, phenotypic, immune, and clinical dimensions, without corresponding to single predefined biological variables. These axes correspond to the dominant sources of variance across the 18-dimensional representation, integrating correlation structure, tumor–normal directionality, survival-associated behavior, microenvironmental gradients, and immune states. By construction, principal components capture orthogonal directions of maximal variance in the joint latent space, providing a data-driven basis for identifying the major phenotypic and regulatory dimensions shaping circuitry geometry.

Dimensions with zero variance were retained but rescaled to unit variance to avoid numerical degeneracies while contributing no effective variance. The first three principal components defined a common three-dimensional latent coordinate system shared by all circuitries and by both sig and int profiles.

Within this three-dimensional PCA space, each circuitry was represented by a pair of local polytopes, one for the sig side and one for the int side. Distances reported throughout the manuscript correspond to Euclidean distances measured in this shared PCA coordinate system. For each 18-dimensional latent vector, we generated 36 vertices by symmetrically perturbing each latent coordinate around its barycenter in an axis-aligned manner. Perturbation magnitudes were proportional to the absolute value of each coordinate, with a minimal ε safeguard to ensure numerical stability. These vertices were projected into the PCA space and used to compute a convex hull for each side, yielding two polytopes per circuitry. In this construction, internal organization reflects the multidimensional contribution structure of latent axes rather than empirical dispersion of individual molecular points. Convex-hull construction and all associated geometric measurements (including volume, anisotropy, and asymmetry) are performed in the full 18-dimensional latent space; three-dimensional convex hulls represent faithful projections of this structure used exclusively for visualization and qualitative inspection.

From this construction, we derived two complementary families of geometric descriptors. First, the barycenter distance between the sig and int polytopes was computed as the Euclidean distance between their three-dimensional centroids, providing a multidimensional measure of regulatory concordance or discordance. Small distances indicate close alignment between regulatory and interacting signatures, whereas larger distances reflect increasing phenotypic and mechanistic divergence across tumor–normal behavior, survival endpoints, microenvironmental features, and immune axes. Second, the volumes of the sig and int convex hulls were quantified as proxies for latent geometric complexity, with small volumes corresponding to low-dimensional, weakly deformed latent profiles and larger volumes indicating high-dimensional, multi-axis deformation consistent with rich, phenotype-coupled circuitry behavior.

To summarize circuitry geometry at scale, we computed for each circuitry: (i) the barycenter distance between sig and int, (ii) the convex hull volume of the sig polytope, (iii) the convex hull volume of the int polytope, and (iv) a volume asymmetry ratio comparing both sides. Using heuristic distance thresholds and data-driven volume quantiles, circuitries were assigned to categorical regimes of distance implication (high concordance, moderate discordance, strong discordance, and extreme discordance) and volume implication (low-dimensional/flat, intermediate, or high-complexity geometries, further stratified by symmetric or asymmetric volume distributions). Together, these descriptors define a geometric phenotype for each circuitry, distinguishing concordant and discordant regulatory behavior, simple versus complex latent organization, and symmetric versus asymmetric regulatory architectures across the multi-omic, immune, and survival dimensions encoded in the OncoMetabolismGPS framework. All geometric constructions, metric computations, regime assignments, and classifications reported in this study were generated using fixed parameters and deterministic rules, without manual curation, interactive tuning, or post-hoc adjustment of thresholds or assignments.

Detailed mathematical definitions and computational implementations of the geometric circuitry framework are provided in the Supplementary Information.

### 2.3. Interactive visualization of geometric metabolic regulatory circuitries

To enable structured exploration and interpretation of the geometric results generated in this study, we developed the SigPolytope Shiny application (URL: https://sigpolytope.shinyapps.io/geometricatlas/). The application provides access to the 18-dimensional latent representations of regulatory circuitries and their corresponding three-dimensional projections, together with derived geometric descriptors including barycenter distance, convex-hull volumes, and associated implication regimes.

Circuitries can be filtered by cancer type, omic and phenotypic layers, metabolic class, pathway annotation, and metabolic cell-death processes. Individual signature–interaction pairs are visualized as interactive dual three-dimensional polytopes within a shared latent coordinate system. In parallel, the interface generates structured textual summaries derived from the standardized signature nomenclature, explicitly linking geometric properties to biological, clinical, microenvironmental, and immune features encoded in the OncoMetabolismGPS framework.

The Shiny application functions as a reproducible visualization and interrogation layer within the analytical workflow, enabling systematic examination of the geometric organization of regulatory circuitries without altering the underlying computational results. All geometric metrics reported in the Results are deterministically derived from the latent tensors and are reproducible from Dataset S1 using the provided codebase, without manual intervention or parameter tuning.

### 2.4. Conceptual definitions

To ensure conceptual clarity and terminological consistency, we introduce here the core geometric constructs that underlie the SigPolytope framework. These definitions formalize how omic signatures and regulatory interactions are represented, compared, and interpreted in latent space, and they establish a common vocabulary for the geometric metrics reported throughout the manuscript. The key terms and measures used in all subsequent analyses are summarized in Box 1.

## 3. Results

### 3.1. Geometric Embedding of 24,796 Regulatory Circuitries in an 18-Dimensional Latent Space

We constructed an 18-dimensional latent tensor for each signature–interaction pair and embedded all circuitries into a unified geometric representation using principal component analysis (PCA). All regulatory circuitries were embedded in a shared 18-dimensional latent space and are visualized throughout the manuscript via a common three-dimensional PCA projection derived from this space. This embedding integrates phenotypic correlation structure, tumor–normal directionality, four survival endpoints, microenvironmental scores, and immune classification into a shared latent representation. The resulting space is continuous but strongly anisotropic, indicating that regulatory circuitries occupy preferential geometric directions rather than forming a homogeneous cloud. Barycenter distances between signature and interaction components follow a long-tailed distribution, with a median of 2.59 (IQR: 1.38–3.99) and extending up to 33.2 (Dataset S1, and Dataset S1 variable descriptors in Supplementary Information), revealing the coexistence of tightly coupled circuitries and profoundly discordant regulatory–metabolic relationships.

Figure 3 (Figure 3_HTML) provides a mechanistic exemplar of this latent organization, illustrating how extreme geometric separation emerges from coherent, multi-layer biological opposition within a single circuitry. In this example, regulatory and interaction components associated with the same phenotypic axis (stemness) localize to distant regions of the latent space, reflecting opposing correlation directionality, survival implications, and microenvironmental and immune contexts. The large barycenter distance and symmetric, high-complexity polytope configuration shown in Figure 3 demonstrate how multidimensional biological divergence is encoded as spatial separation in the latent embedding, thereby establishing an interpretable geometric link between regulatory biology and latent-space topology.

**Figure 3.**
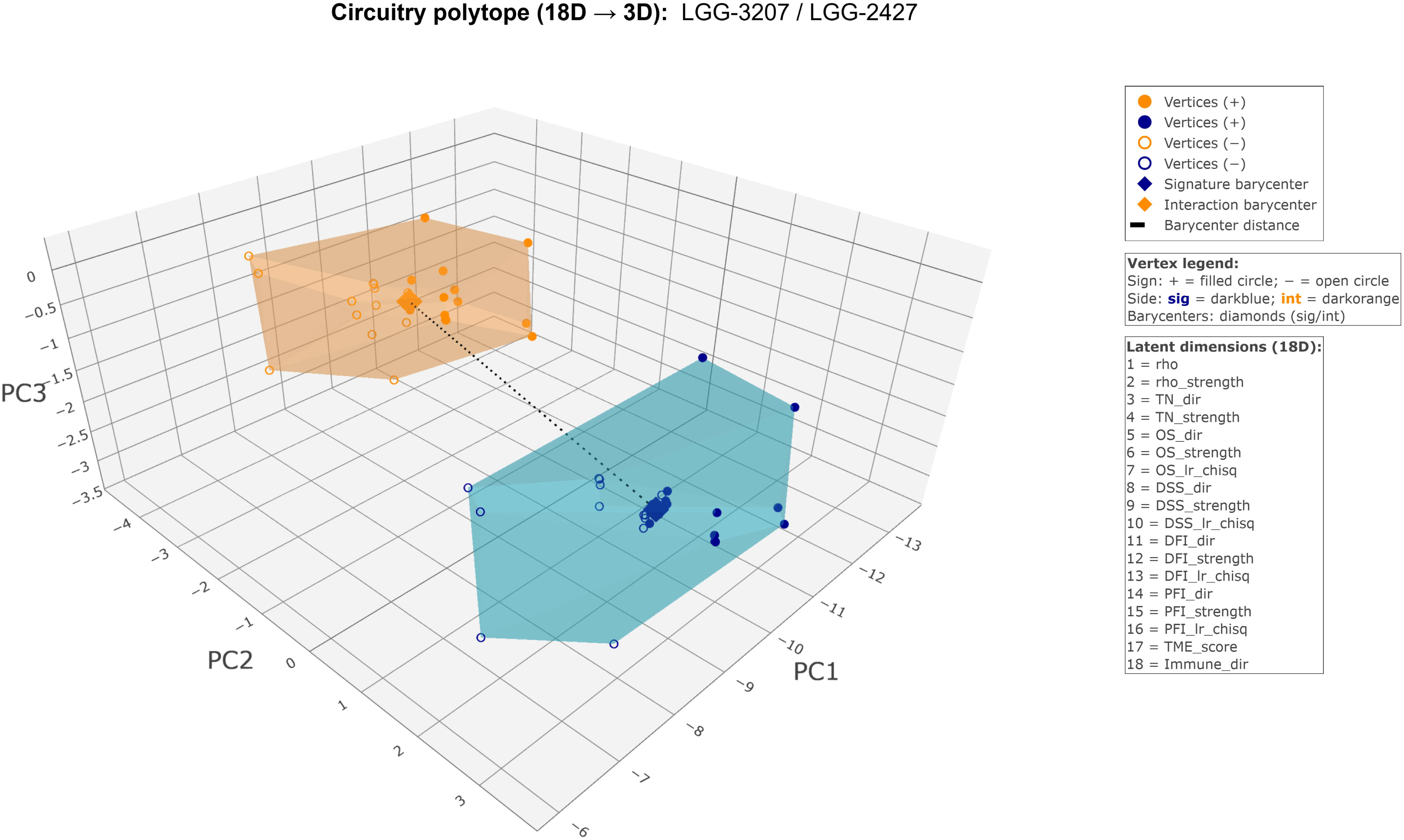
Geometric dual-polytope representation of a highly discordant metabolic regulatory circuitry. Detailed geometric exemplar of a metabolic regulatory circuitry in lower-grade glioma (LGG) linking a miRNA-based regulatory signature (sig: hsa-miR-10a-5p; miRNA expression) and a methylation-derived interaction module (int: CYP8B1; DNA methylation), both associated with the stemness phenotypic layer. The circuitry is embedded within lipid metabolism, specifically the primary bile acid biosynthesis pathway. Although both components map to the same phenotypic domain, they exhibit opposing biological behavior: the regulatory miRNA signature shows a strong negative association with stemness (ρ = −0.41), whereas the metabolic interaction module displays a positive association (ρ = +0.25). This opposition extends to clinical and microenvironmental dimensions, with the sig component associated with a risky survival profile, pro-tumoral microenvironment, and variable immune context, while the int component shows a protective survival association, dual microenvironmental classification, and a cold immune phenotype. Geometrically, the two components are encoded as 18-dimensional latent regulatory vectors and represented as convex dual polytopes in a shared PCA space. The large barycenter distance (7.16) places the circuitry in an extreme discordance regime, while the comparable polytope volumes (symmetric, high-complexity configuration) indicate balanced informational dispersion across layers. Together, these features define an Only Divergent circuitry, exemplifying pronounced geometric and biological dissociation between regulatory and metabolic components within a shared metabolic pathway (Source code and corresponding interactive HTML figure hosted externally in Supporting Information).

### 3.2. Prevalence and Structure of Barycenter-Distance Regimes

Circuitries were assigned to four barycenter-distance (d_bary_) regimes using fixed thresholds in the shared three-dimensional PCA space: high concordance (d_bary_ < 0.5), moderate discordance (0.5 ≤ d_bary_ < 1.5), strong discordance (1.5 ≤ d_bary_ < 2.5), and extreme discordance (d_bary_ ≥ 2.5). These regimes discretize a continuous geometric spectrum and should be interpreted as operational bins rather than intrinsic biological states. This classification summarizes the magnitude of geometric separation between the signature and interaction components of each circuitry in latent space (Table 1A). Importantly, geometric separation does not necessarily imply functional opposition, as spatial divergence in latent space may coexist with concordant behavior along specific annotated biological axes. Because barycenters encode the net directional contribution of all latent dimensions, increasing barycenter distance reflects progressively stronger divergence in overall latent orientation between the regulatory and metabolic sides of a circuitry.

**Table 1.** Distribution of regulatory circuitries across barycenter-distance and volume-based geometric regimes.

**(A) Barycenter-distance regimes**
| **Barycenter-distance implication regime** | **Count**<br>**(n)** | **Percentage**<br>**(%)** |
| --- | --- | --- |
| Extreme discordance | 12869 | 51.9 |
| Moderate discordance | 5360 | 21.6 |
| Strong discordance | 5190 | 20.9 |
| High concordance | 1377 | 5.6 |

**(B) Geometric volume-implication regimes**
| **Geometric volume implication regime** | **Count**<br>**(n)** | **Percentage**<br>**(%)** |
| --- | --- | --- |
| High-complexity multidimensional, strongly asymmetric geometry | 6463 | 26.1 |
| Low-dimensional, strongly asymmetric geometry | 5309 | 21.4 |
| Intermediate-complexity, strongly asymmetric geometry | 3830 | 15.4 |
| Intermediate-complexity, moderately asymmetric geometry | 2836 | 11.4 |
| Low-dimensional, symmetric geometry | 1979 | 8 |
| Intermediate-complexity, symmetric geometry | 1601 | 6.5 |
| High-complexity multidimensional, moderately asymmetric geometry | 1031 | 4.2 |
| Low-dimensional, moderately asymmetric geometry | 978 | 3.9 |
| High-complexity multidimensional, symmetric geometry | 769 | 3.1 |

The distribution of circuitries across barycenter-distance regimes was dominated by discordant geometric classes. Extreme discordance represented the largest regime, accounting for more than half of all circuitries, whereas strong discordance formed a second substantial cluster. Moderate discordance contributed an additional broad intermediate region. High concordance was comparatively rare, comprising only a small fraction of the atlas. These results demonstrate that most circuitries encode divergent latent behaviors in at least one of the major phenotypic axes. Table 1A provides the exact frequencies for all four classes. This pattern establishes discordance as the principal organizing feature of the regulatory landscape, a characteristic readily visible in the separation of barycenters shown in Figure 3.

### 3.3. Convex-Hull Geometry Reveals Nine Classes of Latent Complexity and Asymmetry

Local multidimensional structure around each circuitry was resolved using convex hulls constructed from barycenter-centered perturbations along all 18 latent axes. Hull volume served as a proxy for latent complexity, allowing classification into low-dimensional, intermediate-complexity, and high-complexity geometries. The ratio of signature-to-interaction volumes quantified geometric asymmetry, leading to three possible symmetry states. Together, these measures defined nine mutually exclusive geometric classes.

Across the full Dataset S1, high-complexity multidimensional geometries were prominent, particularly when combined with strong asymmetry between signature and interaction sides. This combination emerged as the single most frequent class, emphasizing that many circuitries engage multiple latent axes yet distribute this complexity unevenly across their two components. Low-dimensional flat geometries were observed but constituted a minority, and the symmetric subset within this tier represented only a modest proportion of the atlas. Intermediate-complexity geometries spanned both symmetric and asymmetric classes and occupied a transitional position in the geometric hierarchy. Table 1B summarizes the distribution of circuitries across all nine structural categories. Figure S1 (<u>Figure S1_HTML</u>) presents representative convex-hull geometries arranged by latent complexity tier (low, intermediate, high) and symmetry state (symmetric vs signature-dominant), providing a visual atlas of the major observed geometric classes.

### 3.4. Joint Patterns of Discordance and Latent Complexity Define Four Principal Geometric Phenotypes

The combination of barycenter-distance regimes with the nine convex-hull categories produced four recurrent geometric phenotypes. Concordant and low-dimensional circuitries formed compact, symmetric dual polytopes with minimal axis deformation. This group was the rarest but represented the most coherent regulatory units in Dataset S1. Concordant but high-dimensional circuitries also emerged, forming large symmetric hulls in which multiple latent axes contributed coordinated deformation on both sides; these circuitries exhibited substantial internal richness while maintaining minimal divergence between their signature and interaction components.

In contrast, discordant low-dimensional circuitries showed pronounced barycenter separation despite minimal hull inflation, indicating sharply opposed yet geometrically simple regulatory programs. The most prevalent phenotype combined discordance with high dimensionality and strong asymmetry. These circuitries displayed large, irregular, and often elongated polytopes with clear separation between signature and interaction barycenters. This dominant class reflects complex and imbalanced phenotypic influences, frequently involving divergent contributions across survival, immune, tumor–normal, and microenvironmental axes.

Figure S2 (<u>Figure S2_HTML</u>) provides population-level exemplars of these four geometric phenotypes. Panel A illustrates a discordant yet low-dimensional asymmetric circuitry, characterized by marked barycenter separation with limited internal complexity. Panel B shows a highly concordant, symmetric, and low-dimensional circuitry, representing compact and internally coherent regulatory organization. Panel C exemplifies extreme discordance with intermediate internal complexity, highlighting phenotypic convergence accompanied by immune and survival divergence. Panel D captures the extreme end of the spectrum, combining maximal discordance with high-dimensional and strongly asymmetric geometry, reflecting deeply imbalanced regulatory and phenotypic influences. Together, these exemplars illustrate how the four principal geometric phenotypes manifest across cancer types and metabolic contexts at the population level.

### 3.5. Representative Circuitries Illustrate Distinct Geometric Behaviors

Detailed visualization of selected circuitries revealed a set of canonical geometric archetypes occupying distinct regions of the joint distance–volume implication space (Figure 4, <u>F4_HTML</u>). Panel A illustrates an extreme discordance archetype in which regulatory and interaction components are widely separated in latent space and form high-complexity, moderately asymmetric dual polytopes, corresponding to pronounced functional divergence between the two sides of the circuitry. By contrast, Panel B exemplifies a high-concordance regime, where regulatory and interaction components localize in close proximity and form symmetric, high-complexity hulls, reflecting coordinated multi-phenotypic integration despite residual immune-level differences. A distinct discordant configuration is shown in Panel C, where large barycenter separation co-occurs with intermediate-complexity, strongly asymmetric hull geometry dominated by the interaction component, capturing a mode of selective divergence in which phenotypic alignment is retained while survival and immune behaviors diverge. Finally, Panel D demonstrates that extreme geometric separation does not necessarily imply biological divergence: despite occupying distant regions of the latent space and forming high-complexity, moderately asymmetric polytopes, both components converge across survival, microenvironmental, and immune dimensions, yielding an Only Convergent circuitry. Together, Panels A–D delineate a coherent set of visually interpretable archetypes, showing how combinations of barycenter distance and polytope volume encode distinct modes of regulatory concordance, divergence, and convergence within the latent embedding.

**Figure 4.**
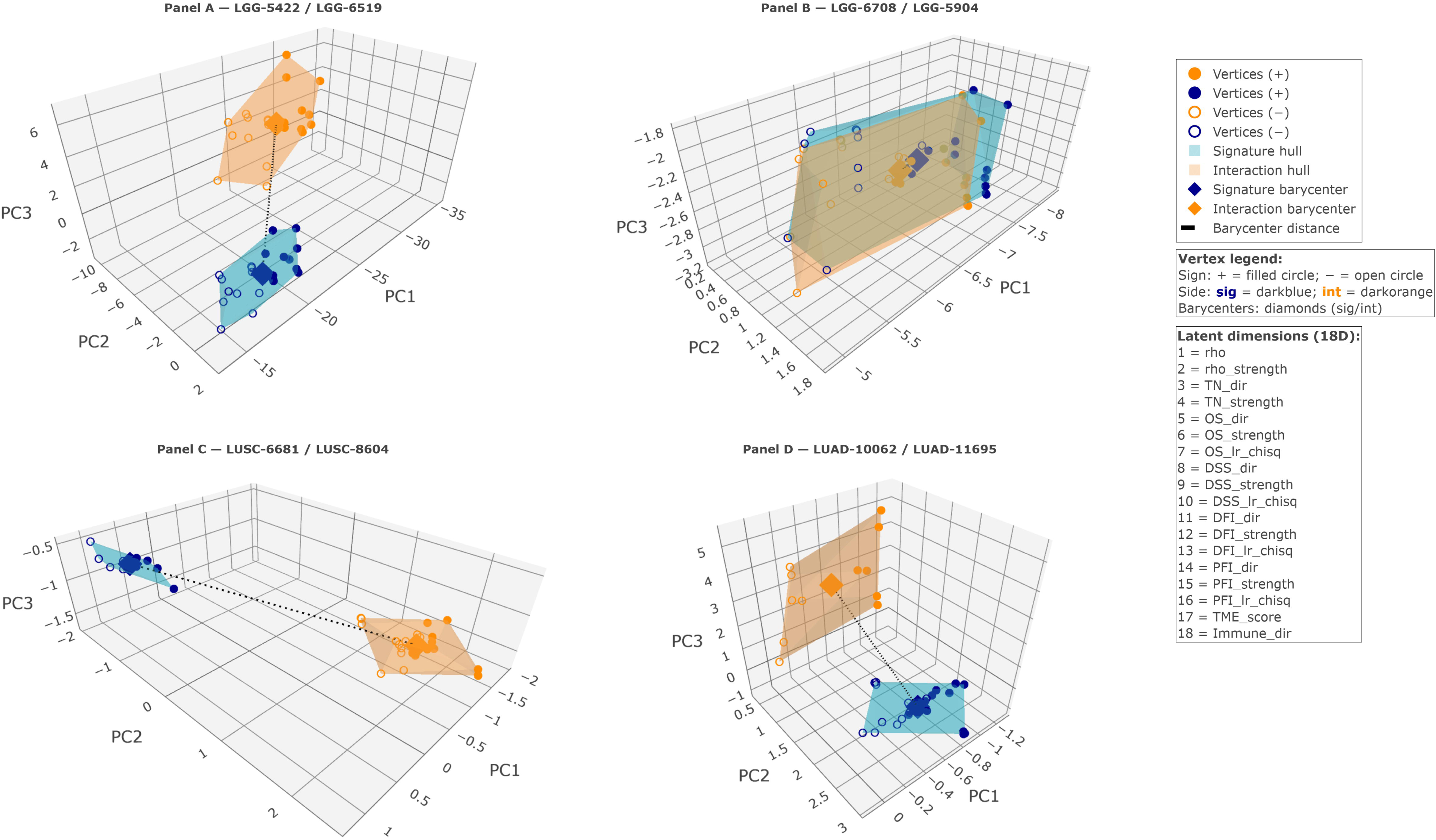
Canonical, visually interpretable archetypes of the joint distance–volume implication space. Panels A–D are ordered to reflect increasing biological resolution rather than increasing geometric distance. (A) Dual-polytope representation of a metabolic regulatory circuitry in lower-grade glioma (LGG) linking a miRNA-based regulatory signature (sig: hsa-miR-615-3p; miRNA expression; stemness) and a CNV-derived interaction module (int: PSAP; CNV; stemness), anchored to lipid metabolism, specifically the sphingolipid metabolism pathway. The two components are widely separated in latent space (barycenter distance = 13.67), defining an extreme discordance regime with high complexity, moderately asymmetric hull geometry. Biologically, the sig component is risky, pro-tumoral, and immune-variable, whereas the int component is protective, associated with a dual microenvironment and a cold immune state, together defining an Only Divergent circuitry. (B) A dual-polytope embedding of a metabolic regulatory circuitry in LGG, coupling a regulatory signature (sig) and an interaction module (int) that are highly aligned across molecular and phenotypic dimensions. The regulatory component corresponds to the lncRNA SNHG12, derived from CNV and associated with stemness, while the interaction component corresponds to KMT2B, likewise CNV-derived and mapped to the stemness phenotypic layer. This circuitry is situated within amino acid metabolism, specifically the lysine degradation pathway. In the shared latent space, the two components localize in close proximity (barycenter distance = 0.20), consistent with a high-concordance geometric regime and a high-complexity, symmetric hull configuration. Biologically, both sig and int components display meaningful risky survival associations and a pro-tumoral microenvironment, while differing at the immune level (variable versus hot), yielding a circuitry characterized by phenotypic and survival convergence with immune divergence. (C) A dual-polytope representation of a metabolic regulatory circuitry in LUSC contrasting a miRNA-based regulatory signature (sig) with a transcriptionally defined interaction module (int) that diverge despite shared phenotypic grounding. The regulatory component corresponds to hsa-miR-30b-5p, derived from miRNA expression and associated with stemness, whereas the interaction component corresponds to LPCAT1, derived from gene expression and likewise mapped to the stemness phenotypic layer. This circuitry is embedded within lipid metabolism, specifically the ether lipid metabolism pathway. In latent space, the two components occupy clearly separated regions (barycenter distance = 3.63), consistent with an extreme discordance regime and an intermediate-complexity, strongly asymmetric hull configuration dominated by the interaction component. Biologically, the sig module displays a meaningful protective survival association together with a pro-tumoral microenvironment and variable immune context, whereas the int module is meaningfully risky, associated with an anti-tumoral microenvironment and a hot immune state, yielding a circuitry characterized by phenotypic convergence with divergent survival and immune behavior. (D) Dual-polytope depiction of a metabolic regulatory circuitry in LUAD juxtaposing a miRNA-based regulatory signature (sig) and a CNV-derived interaction module (int) that diverge geometrically while converging biologically. The regulatory component corresponds to hsa-miR-550a-5p, derived from miRNA expression and associated with microsatellite instability, whereas the interaction component corresponds to NT5C1B, derived from CNV and linked to tumor mutational burden. This circuitry is anchored to nucleotide metabolism, specifically the purine metabolism pathway, in the LUAD context. In the latent embedding, the two components occupy distant regions (barycenter distance = 4.64), consistent with an extreme discordance regime and a high-complexity, moderately asymmetric hull configuration. Biologically, both sig and int components exhibit meaningful risky survival associations, align with an anti-tumoral microenvironment, and share a cold immune phenotype, collectively defining an Only Convergent circuitry despite pronounced geometric separation. (Source code and link to the corresponding HTML figure are in Supporting Information).

### 3.6. Geometric Interpretation and Implications for Regulatory Phenotypes

Barycenter distance provides a direct numerical estimate of the geometric separation between the signature and interaction components of each circuitry. This geometric separation may coexist with functional alignment along specific biological axes. We refer to this pattern as separation-with-convergence, describing circuitries in which the sig and int components are spatially separated in latent space (large barycenter distance) yet exhibit concordant sign structure along specific annotated axes, including survival, microenvironmental, and immune dimensions. In such cases, barycenter distance captures latent-direction separation, whereas the concordance summary class reflects axis-specific alignment, allowing spatial separation to coexist with functional convergence in specific configurations, as illustrated in Figure 4D. In contrast, convex-hull volume quantifies the breadth of latent engagement, indexing how many phenotypic axes contribute to each side. The symmetry measure provides an additional layer of interpretability by identifying unequal contributions to latent complexity.

Together, these measures provide a unified quantitative description of regulatory architecture. Discordant circuitries represent divergent programs whose signature and interaction components occupy distinct regions of latent space. High-complexity geometries indicate multi-phenotype entanglement, whereas low-complexity geometries reflect narrowly tuned regulatory behavior. The predominance of discordant, high-complexity, asymmetric circuitries underscores the structural heterogeneity and phenotypic tension inherent to the regulatory systems captured in Dataset S1. These geometric findings provide a principled, quantitative foundation for subsequent biological analyses that examine how external annotations, such as metabolic pathways, distribute across the landscape defined here.

### 3.7. Distribution of Metabolic Superfamilies Across Geometric Phenotypes

In the analyses that follow, stratification of metabolic superfamilies and pathways is interpreted primarily as a validation of the representational capacity and discriminatory resolution of the geometric framework, rather than as a definitive biological taxonomy of metabolic regulation.

#### 3.7.1. Stratification of metabolic superfamilies across barycenter-distance regimes

The geometric discordance classification derived from barycenter distances revealed consistent large-scale stratification across metabolic superfamilies. All seven classes showed the full range of geometric regimes—high concordance, moderate discordance, strong discordance, and extreme discordance—but with markedly different internal proportions (Figure 5, Dataset S1).

**Figure 5.**
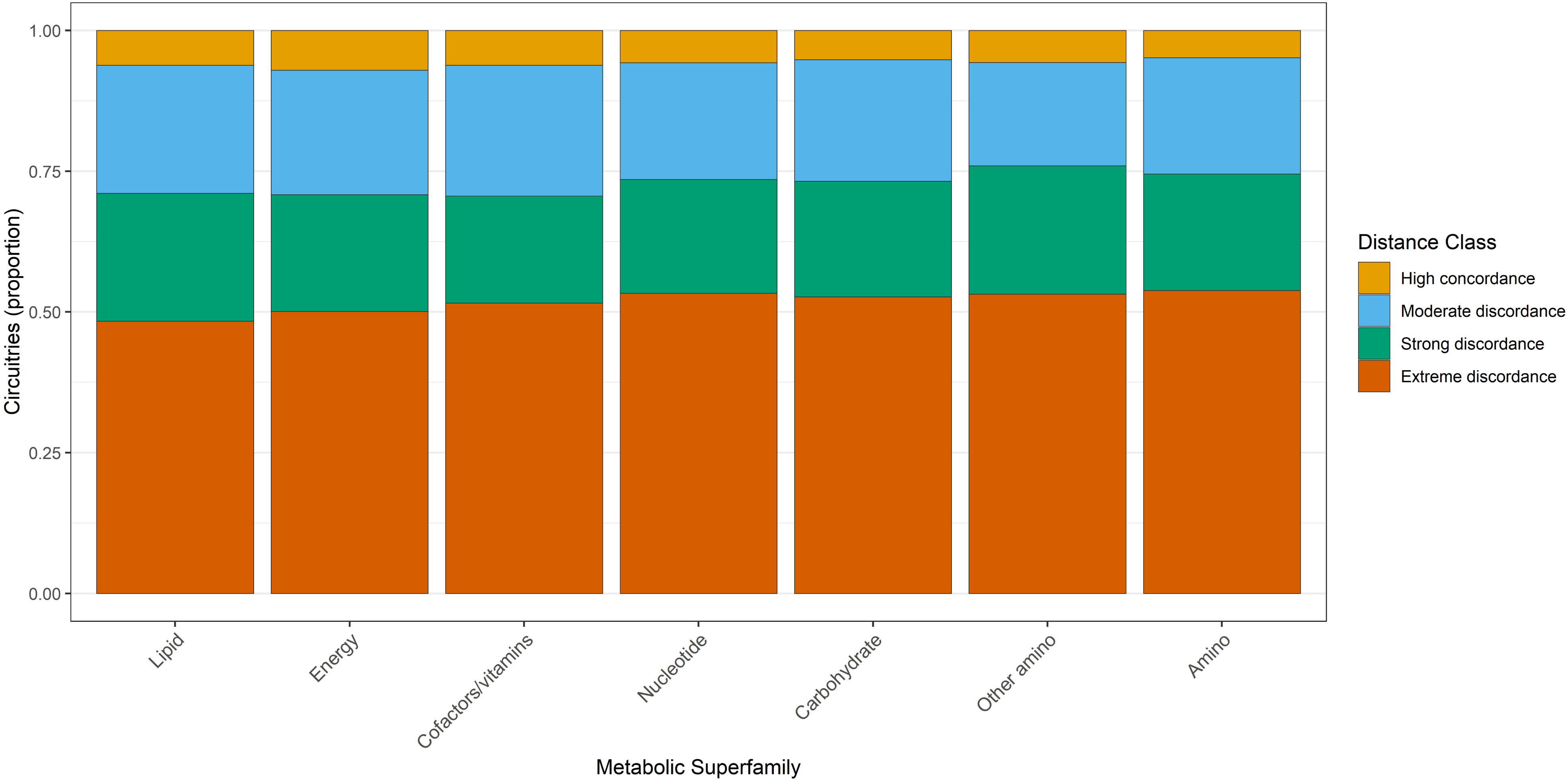
Proportional distribution of regulatory circuitries across geometric distance– implication classes and metabolic superfamilies. Bars represent the fraction of regulatory circuitries assigned to each of the four geometric distance–implication regimes within each metabolic superfamily. Metabolic superfamilies are defined according to KEGG classification and are ordered by Shannon entropy, such that superfamilies exhibiting broader dispersion across geometric regimes appear on the left. This entropy-based ordering highlights differences in geometric organization across metabolic domains: some superfamilies display highly concentrated geometric profiles, indicative of constrained or specialized regulatory behavior, whereas others exhibit more dispersed patterns, consistent with context-flexible or heterogeneous regulatory organization across the circuitry landscape.

Amino acid metabolism exhibited the most polarized distribution, with extreme discordance representing 53.8% of all circuitries assigned to this category (3,491 of 6,489), whereas high-concordance patterns constituted only 4.8% (313 circuitries). Carbohydrate metabolism followed a highly similar structure, with 52.7% of circuitries falling into extreme discordance and 5.2% into high concordance. Energy metabolism demonstrated a modest shift toward more concordant behavior, with 7.0% of circuitries in the high-concordance regime—the largest fraction among the superfamilies—yet the extreme-discordance class still dominated (48.0%).

Lipid metabolism and nucleotide metabolism displayed intermediate distributions, each maintaining ∼50% representation in the extreme-discordance regime (49.8% and 48.7%, respectively). Metabolism of cofactors and vitamins maintained the closest proportional balance between strong and extreme discordance (20.2% and 51.7%, respectively), but still showed only 6.2% high concordance. Across all metabolic classes, moderate and strong discordance each contributed approximately 20% of circuitries, indicating that most superfamilies occupy primarily discordant geometric regimes rather than uniformly concordant ones.

Collectively, these results demonstrate that geometric discordance is not uniformly distributed across metabolic superfamilies; rather, each superfamily exhibits a reproducible internal structure, with extreme discordance consistently dominating but with measurable differences in the magnitude of the high-concordance fraction.

To resolve this organization at finer biological resolution, we next examined barycenter-distance regimes across individual metabolic pathways. Pathway-level stratification revealed coherent gradients of geometric behavior, ranging from pathways dominated by constrained, concordant configurations to those exhibiting broad dispersion across discordance regimes, indicative of pathway-specific regulatory divergence. This distribution is summarized in Figure 6, with pathways ordered by Shannon entropy to highlight differences in geometric pleiotropy.

**Figure 6.**
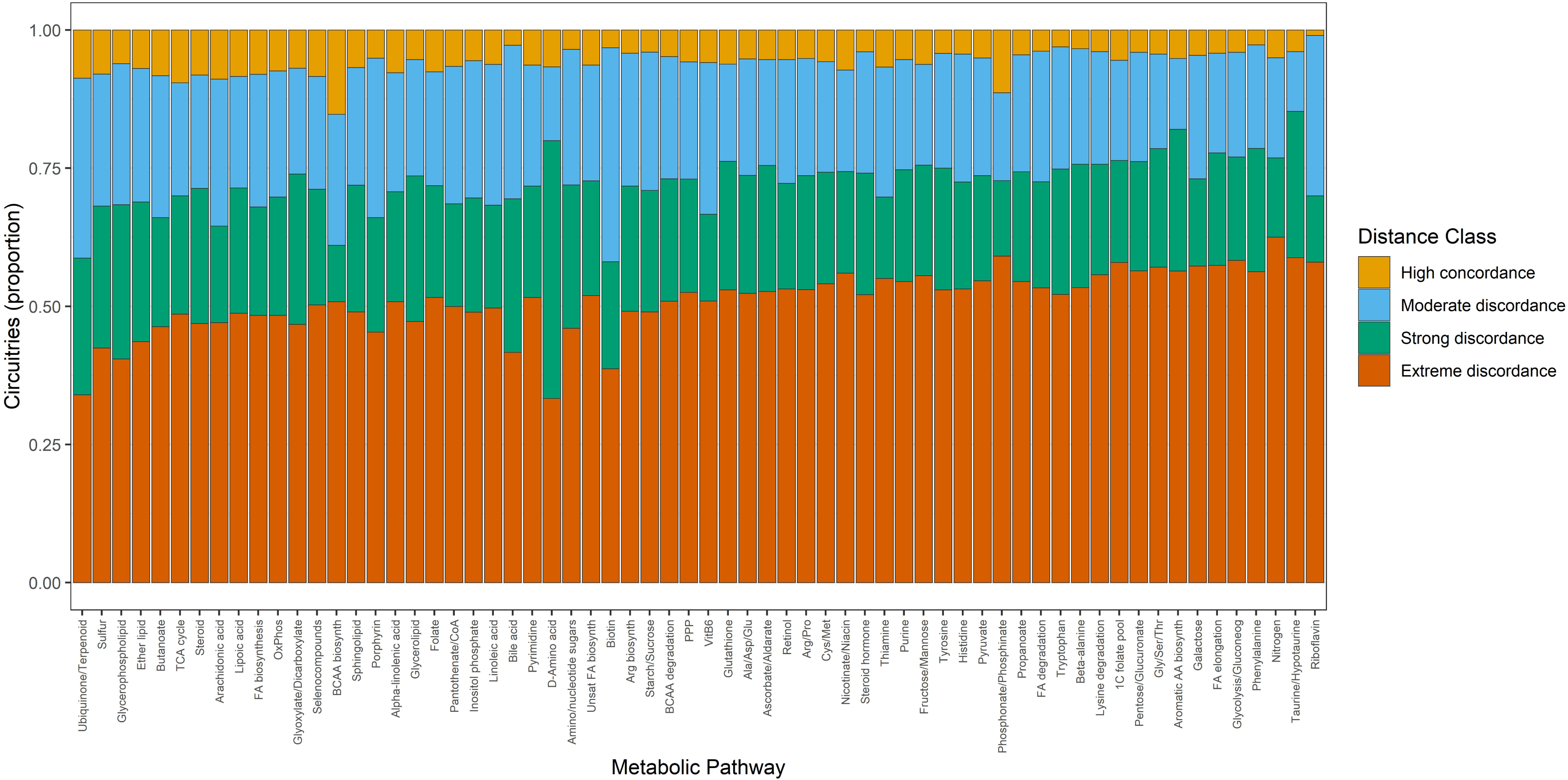
Proportional distribution of regulatory circuitries by geometric distance-implication classes across metabolic pathways. Each bar represents the proportion of circuitries assigned to one of four geometric distance regimes (High concordance, Moderate discordance, Strong discordance, Extreme discordance) within a given metabolic pathway. Pathways are ordered from left to right by Shannon entropy computed over the distance-class distribution, such that pathways exhibiting greater geometric pleiotropy (broad dispersion across discordance regimes) appear earlier. This representation highlights coherent gradients of geometric behavior across metabolic programs, distinguishing pathways that occupy highly constrained geometric states from those displaying structurally diverse circuitry configurations.

#### 3.7.2. Geometric volume implications across metabolic superfamilies

Analysis of convex-hull implications revealed a heterogeneous distribution of latent-space complexity among metabolic superfamilies. All categories included circuitries spanning low-dimensional, intermediate-complexity, and high-complexity regimes, but the relative proportions differed substantially. These superfamily-level distributions of convex-hull–based geometric volume regimes are summarized in Figure 7 (Dataset S1).

**Figure 7.**
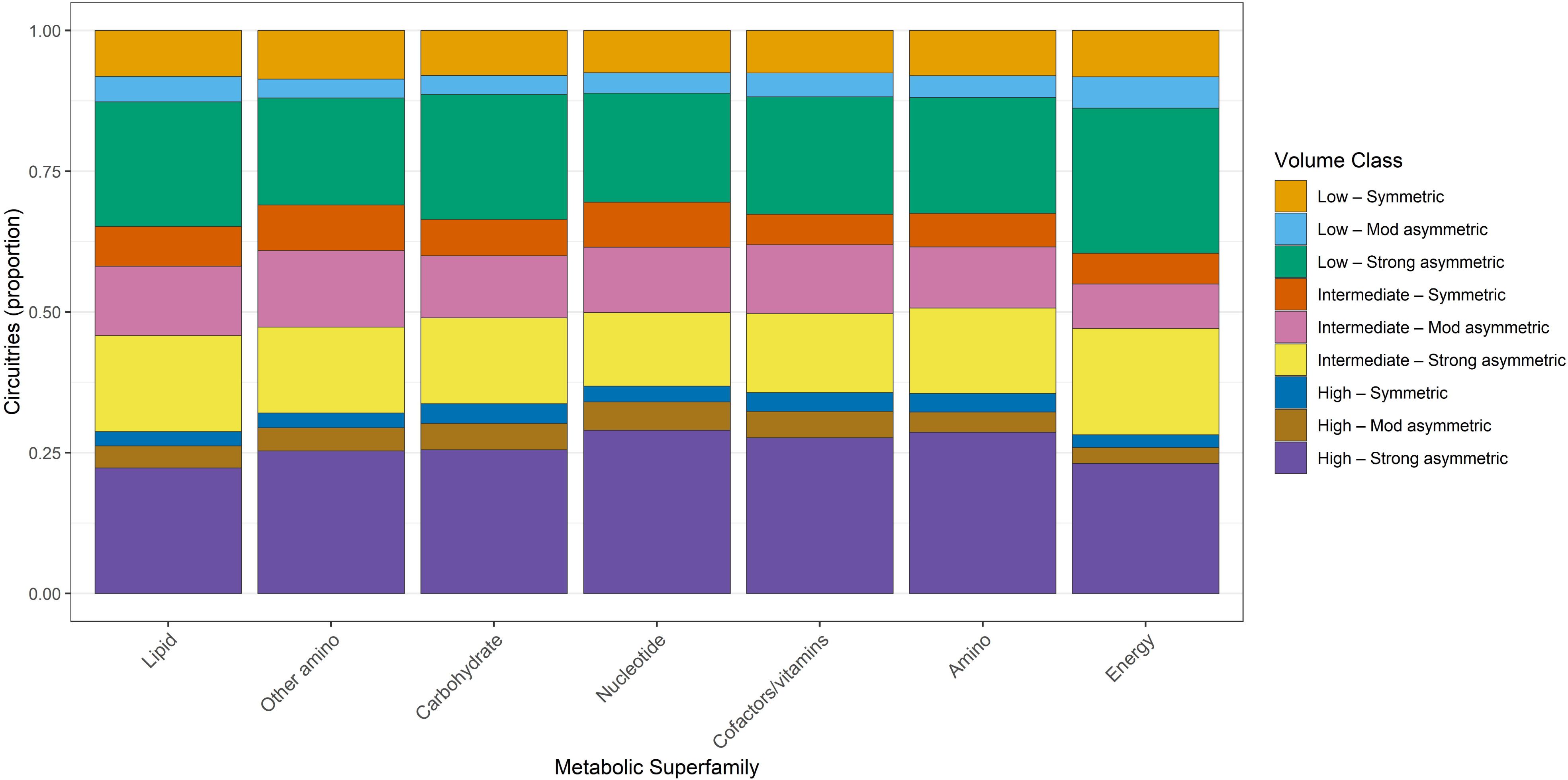
Proportional distribution of regulatory circuitries across geometric volume-implication classes for metabolic superfamilies. Each bar depicts the relative frequency of nine convex-hull–based geometric regimes, defined by latent dimensionality (Low, Intermediate, High) and symmetry/asymmetry of the circuitry volume. Entropy ordering positions superfamilies with high geometric complexity and broad multi-regime occupation to the left, and superfamilies with more restricted geometric signatures to the right. This visualization exposes systematic differences in latent geometric complexity across major metabolic domains.

Amino acid metabolism contained the highest absolute number of high-complexity geometry circuitries, with 44.3% falling into high-complexity multidimensional symmetric regimes and another 30.4% in high-complexity strongly asymmetric configurations. Low-dimensional flat geometries were comparatively rare (collectively 8.8%), indicating that this metabolic class is predominantly associated with higher-order latent deformation patterns.

Carbohydrate metabolism followed a near-identical pattern: 42.8% of circuitries aligned with high-complexity symmetric hulls, while 30.7% mapped to high-complexity strongly asymmetric geometries. Low-dimensional classes again represented <10% of the total. Lipid metabolism showed a similar hierarchical structure, with 43.7% of circuitries in high-complexity symmetric geometries and 28.7% in high-complexity strongly asymmetric geometries.

Energy metabolism exhibited a modestly higher proportion of low-dimensional flat geometries (12.2%) compared with other superfamilies, yet high-complexity regimes remained overwhelmingly dominant (symmetric: 40.3%, strongly asymmetric: 27.9%). Nucleotide metabolism and metabolism of cofactors and vitamins similarly exhibited high-complexity geometries as their prevailing configuration, each with >70% of circuitries occupying high-complexity symmetric or asymmetric regimes.

Taken together, these geometric volume analyses establish that most metabolic superfamilies are preferentially associated with complex latent geometries rather than low-dimensional configurations. Although the degree of asymmetry varies across categories, high-complexity symmetric hulls consistently emerge as the principal geometric phenotype across metabolic domains.

Extending this analysis to the pathway level revealed substantial heterogeneity in latent geometric complexity among individual metabolic programs. While some pathways were concentrated within a limited subset of volume-implication regimes, others spanned the full range of latent dimensionality and convex-hull asymmetry, consistent with pathway-specific multivariate engagement. This fine-grained distribution of volume-implication classes across metabolic pathways is shown in Figure 8, with entropy-based ordering emphasizing differences in geometric specialization versus pleiotropy.

**Figure 8.**
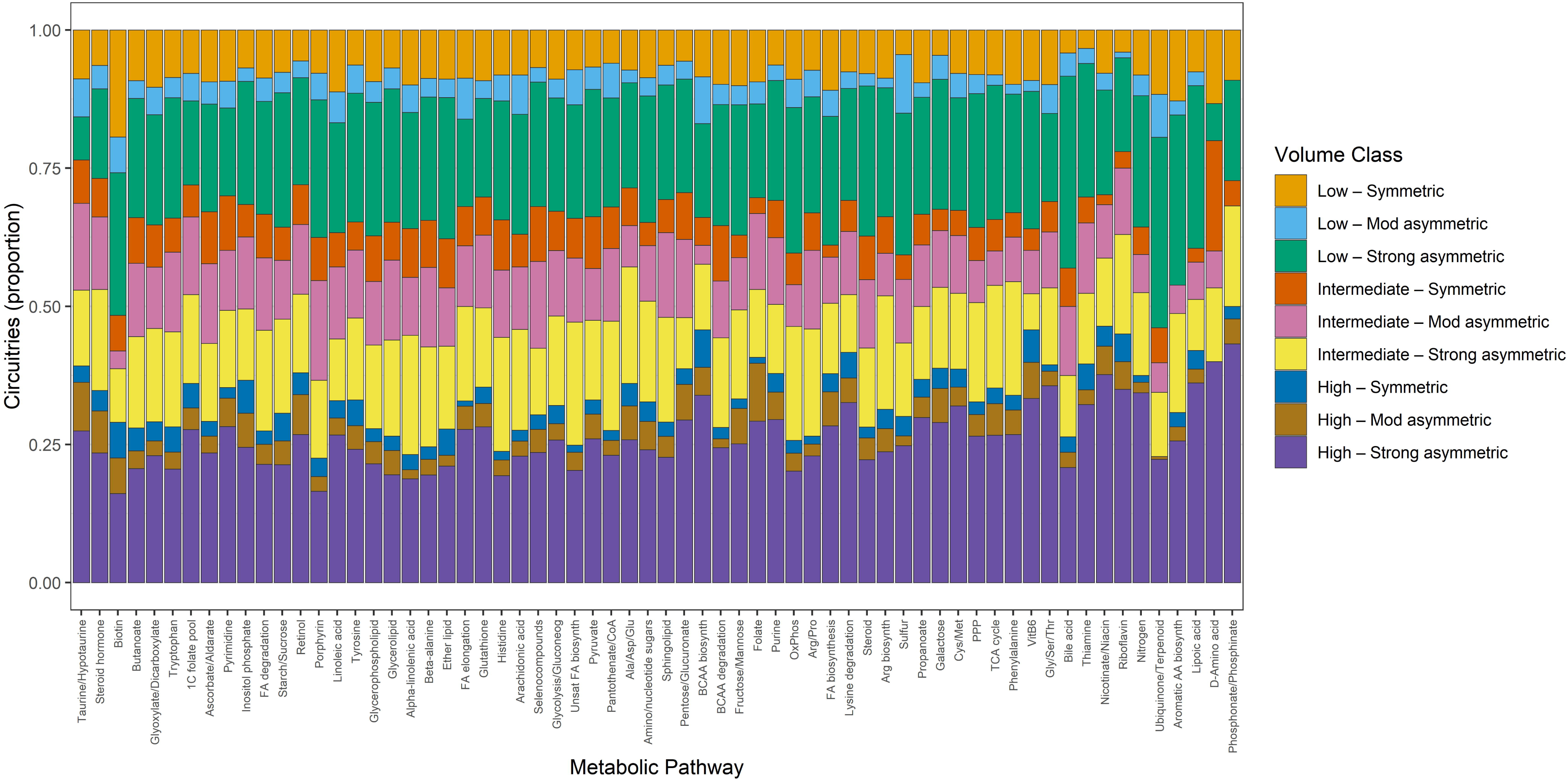
Proportional distribution of regulatory circuitries across geometric volume-implication classes for individual metabolic pathways. Bars represent the distribution across nine geometric volume regimes, capturing both latent dimensionality and convex-hull asymmetry. Pathways are ordered by Shannon entropy computed from the volume-class distribution, revealing a continuum from highly pleiotropic pathways that participate in diverse geometric architectures to specialized pathways concentrated in one or a few geometric regimes. The figure provides a fine-grained map of geometric heterogeneity across metabolic circuitry structures.

Across both geometric dimensions—discordance magnitude (distance) and latent complexity (hull volume)—the metabolic superfamilies exhibit reproducible, non-uniform distributions. Extreme discordance dominates every metabolic class, indicating that signatures and interactions typically encode divergent latent directions. Simultaneously, high-complexity multidimensional geometries represent the major structural phenotype within each metabolic superfamily, supporting the interpretation that metabolic regulatory interactions generally recruit multiple latent axes rather than being confined to narrow or low-dimensional modes.

### 3.8. Entropy-Based Quantification of Geometric Heterogeneity Across Concordance Classes

We next examined how circuitries distribute across the full spectrum of multivariate concordance outcomes. By quantifying the proportion of circuitries assigned to each final concordance summary class and ordering these categories by Shannon entropy, we visualized the relative structural diversity underlying each outcome state (Figure S3). The entropy-based ordering provides a neutral ranking that places classes with broader geometric dispersion earlier in the plot, facilitating comparison of distributional patterns across the entire concordance landscape. In this context, entropy captures the dispersion of circuitries across the four geometric regimes (high concordance, moderate discordance, strong discordance, and extreme discordance). Because these regimes encode distinct latent structural configurations of the underlying multi-omic circuitry, entropy provides a principled measure of whether biologically defined categories occupy a narrow versus diffuse region of the geometric space. Low entropy reflects structural concentration, indicating that circuitries within a concordance class converge toward a dominant geometric configuration. High entropy, in contrast, denotes a broad distribution across geometric regimes, implying that circuitries with similar biological outcomes may arise from widely varying multivariate architectures.

Applying this framework revealed a nontrivial decoupling between biological concordance and geometric organization. The classes Only Convergent and Only Divergent, which together represent the two largest categories in Dataset S1, exhibited markedly different entropy values despite their comparable sample sizes. Only Convergent displayed a higher entropy (∼1.30), indicating that these circuitries populate a broad range of geometric regimes without a single dominant configuration. This suggests that fully concordant multi-omic signals can emerge from diverse structural organizations, reflecting multiple mechanistic routes to geometric agreement. By contrast, only Divergent exhibited lower entropy (∼1.07), demonstrating a more concentrated geometric profile in which divergent biological signals disproportionately arise from a narrower subset of geometric configurations. This asymmetry implies that divergence in immune, phenotype, Cox, or survival dimensions may be driven by specific, structurally constrained geometric patterns, whereas full concordance is structurally more permissive.

Together, these findings demonstrate that entropy provides a sensitive discriminator of geometric heterogeneity across concordance categories and uncovers an emergent organizing principle: biological divergence tends to be geometrically constrained, whereas biological concordance is realized through a richer diversity of multivariate configurations. This insight strengthens the mechanistic interpretability of the geometric framework and supports the notion that distinct biological agreement states arise from characteristic, and quantifiable, geometric architectures.

### 3.9. Shiny-based interpretation of geometric signatures

Interactive exploration of OncoMetabolism regulatory circuitry geometry was performed using the SigPolytope Shiny application, which enables direct visualization and comparison of convex-polytope representations of multi-omic regulatory circuitries. This application provides a practical interpretive interface for the geometric structures described in this atlas. Through navigation of the shared latent space, users can examine how circuitries distribute across regimes of concordance and discordance, levels of geometric complexity, and degrees of asymmetry between signature and interaction components.

Joint visualization of dual polytopes together with automatically generated textual summaries allows abstract geometric properties—such as barycenter displacement and multidimensional expansion—to be systematically linked to biological and clinical attributes, including survival associations, tumor microenvironmental context, and immune phenotypes. This interactive framework supports the identification of representative circuitries, comparison of structurally related signatures, and detection of regulatory divergences that are not readily apparent from conventional vector-based or network-based representations.

## 4. Discussion

### 4.1. Rethinking How Multi-Omic Regulatory Interactions Are Represented

This study is methodological and conceptual in scope: it introduces a geometric framework for representing and comparing multi-omic signatures and regulatory circuitries, rather than extending or replacing existing biological atlases or making direct causal claims about specific molecular mechanisms. The multidimensional structure revealed through this framework necessitates a fundamental reconsideration of how multi-omic interactions are represented, visualized, and interpreted. Multi-omic circuitries are not linear associations; they are high-dimensional informational entities whose behavior arises from coordinated contributions across molecular, phenotypic, immune, microenvironmental, and clinical axes. Yet the dominant visual paradigms employed in the field—network maps, alluvial flows, layered cubes, and force-directed 2D or 3D graphs—compress this complex architecture into schematic representations that obscure structural features essential for mechanistic interpretation. The analyses presented here illustrate that such representational flattening is more than an aesthetic constraint: it acts as a conceptual bottleneck that limits what can be inferred from multi-omic data.

Network diagrams, whether in two or three dimensions, excel at depicting connectivity but lack any meaningful metric geometry. Physical proximity between nodes reflects the behavior of a layout algorithm rather than biological organization. Circuitries that diverge strongly in latent space may appear arbitrarily close if the force-directed algorithm attracts them, while highly similar entities may be placed far apart for reasons unrelated to biology. Networks, therefore, encode who connects to whom, but cannot express how regulatory signals distribute across latent dimensions, nor the structural tensions, concordances, or divergences that emerge from multi-omic integration.

Alluvial diagrams and multi-omic cubes provide alternate perspectives yet remain fundamentally constrained. Alluvial representations capture continuity—how features migrate across categories or pathway memberships—but they lack any ability to depict curvature, anisotropy, or latent-dimensional deformation. Multi-omic cubes restore interpretability along observable axes such as tumor–normal directionality, immune polarization, or survival tendencies, but their fixed coordinate system cannot encode the internal covariance structure that governs circuitry geometry. They cannot reveal asymmetric expansion, multidirectional deformation, or barycenter displacement—structural elements that are central to understanding phenotypic divergence.

The convex-polytope framework introduced here directly addresses these limitations. By embedding each circuitry as a pair of 18-dimensional latent regulatory vectors, expanding them into barycenter-centered point clouds, and projecting them into a shared geometric space, the resulting dual hulls preserve structural information rather than abstracting it away. Hull volume quantifies latent complexity, barycenter distance quantifies discordance, and anisotropy captures directional dominance across phenotypic or mechanistic axes. Crucially, these properties arise intrinsically from the data, not from visualization heuristics or graph-layout algorithms.

This distinction becomes decisive when examining the Dataset S1. More than half of all circuitries fall into extreme-discordance regimes, and more than 70% occupy high-complexity symmetric or strongly asymmetric geometries. No alluvial diagram, cube, or network graph can readily reveal this structural prevalence, nor explain why discordant, high-dimensional asymmetric circuitries dominate metabolic regulatory space. Most directly, the dual-polytope geometry exposes these large-scale organizational principles: how signature and interaction components diverge across latent dimensions, how multi-axis deformation reflects underlying mechanistic heterogeneity, and how regulatory modules distribute systematically across metabolic programs.

Accordingly, geometry is not simply an alternative visualization; it constitutes a representational framework that retains multidimensional structure and enables interpretive statements unavailable through conventional approaches. The convex hull does not merely depict a circuitry—it is the circuitry, expressed as a measurable geometric object whose biological meaning arises from its intrinsic properties rather than imposed graphical conventions. This geometric framing anchors the remainder of the Discussion, offering a rigorous foundation for interpreting phenotypic discordance, latent complexity, and the distribution of metabolic processes across geometric regimes.

Because a hull is defined over a set of points in latent space, it admits intrinsic measurements that translate naturally into biological interpretation. Hull volume reflects multidimensional amplitude, indexing the breadth of mechanistic and phenotypic dispersion. Principal-axis orientation, derived from the covariance structure of latent coordinates, reveals which biological dimensions dominate a circuitry’s behavior—whether regulation is driven by survival, immune directionality, tumor–normal polarity, or microenvironmental influence. Curvature and anisotropy differentiate coherent single-axis programs from modular, multi-axis, or branching architectures.

These geometric metrics formalize patterns that biologists often perceive qualitatively but cannot quantify using list-based or vector-based representations, thereby providing a robust bridge between biological intuition and computational structure. Importantly, geometric formalism does not replace biological reasoning; it strengthens it by making structural relationships explicit. A convex hull can show that a seemingly unified signature is in fact topologically fragmented, prompting mechanistic subdivision. Conversely, it may reveal that two signatures with distinct molecular compositions are geometrically superimposable, implying functional equivalence and potential biomarker redundancy (Figure 4B, <u>F4_HTML</u>; circuitry_id = “LGG-6708 / LGG-5904”).

Through this lens, core biological concepts—heterogeneity, redundancy, convergence, divergence, modularity, dominance—cease to be metaphors and become quantifiable geometric properties. By applying geometric reasoning to multi-omic signatures, analysis acquires explanatory power not possible with traditional representations. The convex hull thus becomes not merely a visualization artifact, but an epistemological bridge linking multi-omic evidence to biological meaning.

### 4.2. Geometry as a Foundation for Signature Comparison, Classification, and Translational Relevance

The geometric framework introduced above reorients the analysis of multi-omic signatures from describing each signature in isolation to understanding how signatures relate to one another as structured informational entities. Once signatures are embedded in a common latent coordinate system and expressed as convex polytopes, similarity, divergence, redundancy, complementarity, and mechanistic distance become measurable quantities rather than qualitative impressions. Geometry thus shifts the interpretive focus from enumeration to comparison and from annotation to structure.

Two signatures that occupy overlapping hull volumes can be interpreted as mechanistically aligned—even if they share few or no molecular components—because geometric overlap reflects concordant behavior across the 18 latent axes that encode correlation structure, tumor–normal shifts, survival tendencies, microenvironmental influence, and immune state (Figure 4). Conversely, signatures that diverge markedly in orientation, volume, or curvature occupy distinct regions of latent space and therefore encode distinct biological programs (Figure 3). Through this lens, geometric representation transforms signature analysis from a descriptive exercise into a comparative science of multidimensional biological architectures.

Because geometric relationships encode concordance, divergence, and redundancy across clinically relevant latent dimensions, this comparative capacity has immediate translational consequences. In precision oncology, numerous candidate signatures are often proposed for the same phenotype, pathway, or clinical endpoint. Lacking a structural framework, these signatures are typically treated as independent entities, leaving redundancy, equivalence, or complementarity to conjecture rather than demonstration. The convex-polytope representation resolves this ambiguity: redundancy appears as volumetric overlap, equivalence as geometric congruence, and complementarity as divergence along orthogonal axes. Geometry, therefore, enables principled prioritization—redundant signatures can be collapsed, divergent signatures can be stratified, and only the most structurally informative candidates should advance to clinical validation.

The interpretive power of geometric reasoning becomes particularly clear in immuno-oncology, where immune states define well-characterized but multidimensionally coupled phenotypic regimes. Immune-active signatures that converge on similar latent phenotypes produce hulls that occupy overlapping regions of geometric space (Figure 4), while immune-excluded or immune-suppressed signatures diverge along axes associated with stromal shielding, metabolic reprogramming, or immune evasion (Figure 3). This stratification is not imposed; it emerges directly from the internal geometry of each circuitry. In this context, geometry becomes a functional classifier: hot-aligned geometries identify candidate predictors of immunotherapy benefit, whereas cold-aligned geometries reveal signatures associated with immune exclusion, immunoediting trajectories, or therapeutic resistance.

Beyond immune phenotypes, geometric divergence offers a rational framework for therapeutic inference. Candidate drugs, synthetic-lethality hypotheses, and combination strategies can be prioritized based on mechanistic distance—defined geometrically—rather than superficial gene overlap or heuristic similarity scores. Signatures separated by large barycenter distances represent mechanistically discordant programs that may respond to orthogonal therapeutic pressures. Signatures with minimal barycenter separation and congruent volumes, by contrast, highlight biological redundancies that should be avoided when constructing multi-signature panels.

More broadly, geometric representation provides a foundation for de-redundancy, benchmarking, and cohort stratification across oncology applications. Two prognostic or predictive signatures purported to stratify the same patient population can be compared directly: if their hulls are congruent, only one is necessary; if they diverge, each captures orthogonal axes of tumor biology, justifying complementary use. Geometry thus transforms signatures from narrative artifacts into quantitative biomarker objects—entities that can be ranked, merged, or discarded based on structural justification rather than interpretive intuition.

Although barycenter distance and hull-volume ratio jointly contribute to discordance classification, these metrics quantify orthogonal aspects of geometry: positional displacement versus latent-dimensional expansion. Their combination accounts for the broad nonlinear behavior observed in extreme-discordance circuitries and underscores the multidimensional nature of regulatory divergence. By giving signatures a geometry, this framework provides a rigorous foundation for comparative reasoning, biologically meaningful classification, and translational prioritization. Geometry distinguishes signal from redundancy, mechanism from noise, and biological architecture from statistical coincidence. It is not an embellishment of visualization but an enabling condition for the next generation of mechanistically grounded, clinically credible biomarker science.

### 4.3. Limitations and Caveats

Although the geometric framework presented here provides a principled method for representing, comparing, and interpreting multi-omic regulatory circuitries, several limitations should be acknowledged. These limitations do not undermine the framework’s core contributions but clarify the boundaries within which the current implementation should be interpreted and extended.

First, the latent space is constructed from an 18-dimensional tensor that captures the dominant measurable axes of circuitry behavior—correlation structure, tumor–normal directionality, survival tendencies, microenvironmental gradients, and immune states. While these variables encompass the major phenotypic dimensions encoded in the OncoMetabolismGPS atlas, they are not exhaustive. Importantly, the conclusions derived here depend on relative geometric organization rather than exhaustive axis coverage. Additional regulatory layers, such as chromatin topology, spatial transcriptomics, metabolic flux measurements, or treatment-induced perturbations, may introduce new axes of variation not captured by the present latent construction. Future versions of the atlas will benefit from integrating these modalities, particularly in contexts where regulatory behavior is governed by non-transcriptomic constraints.

Second, the PCA projection used to obtain a common geometric coordinate system necessarily compresses high-dimensional variation into three dimensions for visualization. This projection preserves dominant variance but cannot represent all latent interactions with equal fidelity. While the convex-polytope operations occur in the full 18-dimensional space before projection, the interpretability of hull shape, curvature, or anisotropy may still be influenced by the geometry of dimensionality reduction. Alternative embeddings (e.g., manifold learning, diffusion geometry, generalized linear factor models) may reveal orthogonal structures not captured by PCA and warrant comparison in future studies. Although the numerical thresholds used to define barycenter-distance regimes are heuristic, they are applied uniformly across all circuitries and yield stable qualitative patterns across cancer types, metabolic classes, and regulatory contexts. These thresholds serve as operational boundaries for summarizing continuous geometric variation rather than as sharp biological cutoffs, and the resulting regime structure is robust to moderate threshold perturbation.

Third, the convex-hull expansion relies on symmetric perturbations around each latent coordinate, allowing hulls to encode multidimensional deformation around a barycenter. Although this perturbation strategy preserves numerical stability and structural interpretability, it represents one of many possible ways to induce local geometric neighborhoods while preserving strict comparability across thousands of circuitries analyzed under identical geometric constraints. More complex or data-adaptive expansions—such as anisotropic perturbation kernels, distribution-based vertex generation, or probabilistic hulls—may refine the geometric representation in settings where circuitry behavior is highly non-linear.

Fourth, the interpretations presented here depend on the accuracy and completeness of underlying omic measurements. Batch effects, missingness patterns, or biases in clinical annotation can influence latent-axis contributions, especially in phenotypes with sparse or noisy endpoints. Although the imputation, normalization, and quality-control steps employed in the OncoMetabolismGPS pipeline adhere to stringent standards, residual artifacts may propagate into latent geometry. Cross-cohort validation, perturbation testing, and sensitivity analyses will be essential for establishing the robustness of the geometric classes across datasets, disease contexts, and platforms.

Fifth, the framework models each circuitry as a static entity derived from bulk omic measurements. This limitation reflects data availability rather than a constraint of the geometric formalism itself. Tumors, however, evolve dynamically, and single-cell heterogeneity, clonal substructure, and microenvironmental flux may generate temporal or spatial geometry shifts. Extending the convex-polytope model to longitudinal datasets or to per-cell circuitry embeddings may reveal regulatory dynamics that remain inaccessible in static representations.

Finally, although geometry captures structural relationships with high fidelity, it does not encode causal directionality. Barycenter displacement, hull inflation, and axis alignment quantify how signatures relate in latent space but do not specify why these relationships arise or which regulatory interactions drive them. Integrating geometric representations with causal graph inference, perturbation experiments, or mechanistic pathway models will be an important next step toward linking structure with mechanistic explanation.

Collectively, these limitations underscore the need for continued methodological refinement while reinforcing the conceptual strength of the geometric approach. The convex-polytope framework establishes a scalable foundation for representing multi-omic regulatory entities, and its limitations highlight natural directions for extending the model into causal, dynamic, and multimodal domains.

### 4.4. Concluding Remarks

This work reconceptualizes omic signatures as multidimensional informational entities whose structure and biological meaning are inherently geometric, rather than vectorial or list-based. By embedding 24,796 metabolic regulatory circuitries into an 18-dimensional latent space and representing each as a dual convex polytope, we establish that signatures possess definable geometric identities—comprising barycenter displacement, latent complexity, anisotropy, and volume asymmetry—that expose mechanistic and phenotypic relationships inaccessible to conventional representational frameworks.

A key insight from the atlas is that discordance, not concordance, dominates the regulatory landscape. Most circuitries fall into strong or extreme discordance regimes and exhibit high-dimensional, frequently asymmetric geometries. These findings overturn the implicit assumption that multi-omic signatures function as coherent, unified objects; instead, they behave as structurally heterogeneous constructs whose biological meaning emerges from multi-axis deformation patterns rather than from molecular membership alone.

By treating each circuitry as a measurable geometric object, this framework enables principled comparison, stratification, and prioritization of signatures. Overlapping hulls reveal functional alignment independent of shared molecular components; divergent hulls quantify mechanistic distance; and symmetric or asymmetric volumes distinguish balanced from imbalanced phenotypic engagement. These geometric relationships support translational decisions— ranging from de-redundancy of biomarker panels to mechanistic interpretation of immune ecology and the rational selection of prognostic or therapeutic signatures.

More broadly, geometric formalism bridges biological intuition with computational structure. Properties traditionally described qualitatively—heterogeneity, modularity, convergence, divergence, dominance—become computable traits embedded in latent space. This transition from narrative to structure defines the conceptual pivot of the present work and establishes geometry as a rigorous language for expressing the organizational principles of multi-omic regulatory systems.

The resulting geometric atlas provides a scalable foundation for future multi-omic discovery. As emerging data modalities—including spatial transcriptomics, chromatin topology, metabolic flux profiling, and longitudinal tumor dynamics—are integrated into latent space, the polytope representation will expand to capture increasingly complex regulatory architectures. This trajectory points toward a new analytic paradigm in which mechanisms, phenotypes, and contexts are represented as geometric objects amenable to comparison, classification, and causal inference.

By giving omic signatures a geometry, we equip multi-omic science with a structural vocabulary capable of capturing the richness, heterogeneity, and emergent organization of cancer biology. The geometric framework and atlas presented here constitute not only an analytical resource but a conceptual foundation for building the next generation of mechanistically grounded, clinically actionable biomarker systems.

## Supporting information

Data variable descriptors

Regulatory circuitries

## Statements

### Data availability statement

All data and R source code generated or analyzed during this study, including processed multi-omic matrices, derived geometric features, and annotated regulatory circuitry tables, are provided in the Supplementary Information and associated Supplementary Tables. The complete computational codebase used to construct the latent representations, geometric embeddings, and figures is publicly available via the SigPolytope repository (https://github.com/BioCancerInformatics/SigPolytope), together with interactive HTML figures used during analysis and quality control. Static, publication-grade figures shown in the manuscript were regenerated deterministically from the underlying data using non-interactive plotting pipelines. Portions of the R code were initially drafted with the assistance of ChatGPT (OpenAI; model version: GPT-5.2) as a programming aid. All scripts were subsequently modified, programmatically audited, and logically validated by the authors using independent verification routines prior to use in the analyses.

### Ethics Statement

No human or animal subjects were directly involved in this study. All analyses were conducted using previously generated, de-identified molecular, clinical, and demographic data obtained from publicly available resources. As such, no ethical approval or informed consent was required for this work.

### Author Contributions

H.A.C.N. contributed to the conceptual development of the study, developed the SigPolytope Shiny applications, and participated in the review of manuscript drafts. E.M.-A. conceived the study, designed the conceptual and methodological framework, supervised the analyses, and wrote the manuscript. All authors reviewed and approved the final version of the manuscript.

## Funding

This work received institutional support from the Programa de Apoio à Pesquisa Institucional (PAPIC; Grant UENF001/2024) and PAPIC PLUS (Grant UENF001/2025), awarded to E.M.-A. H.A.C.N. was supported by a doctoral scholarship from the Fundação de Amparo à Pesquisa do Estado do Rio de Janeiro (FAPERJ) and by the Coordenação de Aperfeiçoamento de Pessoal de Nível Superior (CAPES), Brazil, through the Doctoral Sandwich Program (PDSE -88881.219120/2025-01). Additional infrastructure funding was provided by the Financiadora de Estudos e Projetos (Finep) and the National Fund for Scientific and Technological Development (FNDCT) under the PROINFRA 2021 program (Agreement No. 0.1.22.0442.00).

## Acknowledgments

The authors thank Emanuell Rodrigues da Silva and Victor dos Santos Lopes for helpful discussions during the early stages of the development of this manuscript.

## Conflict of interest

The authors declare that the research was conducted in the absence of any commercial or financial relationships that could be construed as a potential conflict of interest.

## Generative AI statement

Generative artificial intelligence tools were used in a limited and supervised manner during manuscript preparation. ChatGPT (OpenAI; model version: GPT-5.2) was employed to assist with English language revision, grammar checking, and syntactic refinement of selected text passages. ChatGPT was also used as an auxiliary aid during the initial drafting of R code. All AI-assisted code was subsequently edited programmatically by the authors and incorporated into a fully controlled computational pipeline that included explicit auditing, logical validation, and output-consistency checks. The authors take full responsibility for the content of the manuscript, including the accuracy of the text, the correctness of the code, and the validity of all analyses and interpretations.

## Publisher’s note

All claims expressed in this article are solely those of the authors and do not necessarily represent those of their affiliated organizations, or those of the publisher, the editors and the reviewers. Any product that may be evaluated in this article, or claim that may be made by its manufacturer, is not guaranteed or endorsed by the publisher.

---

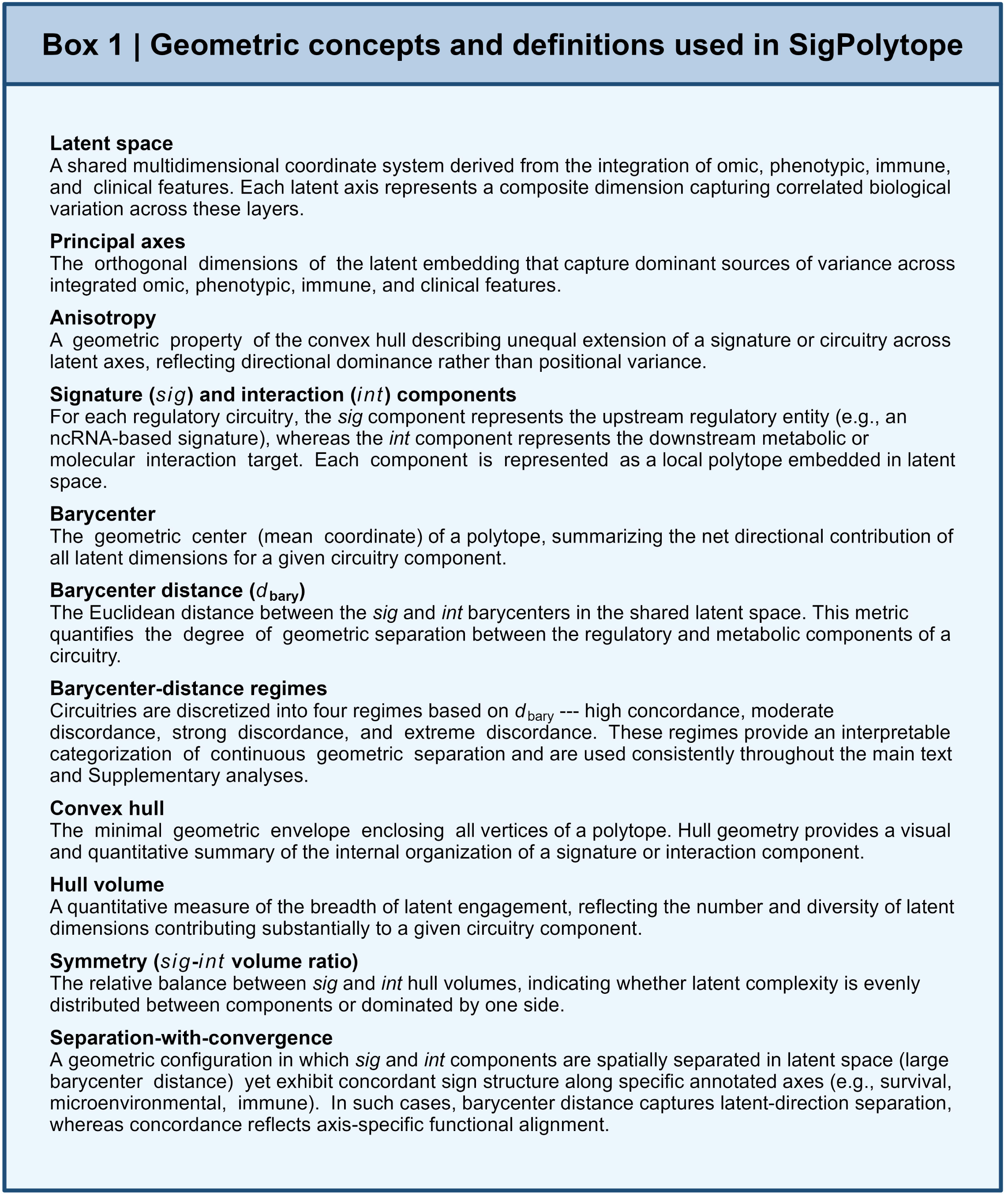

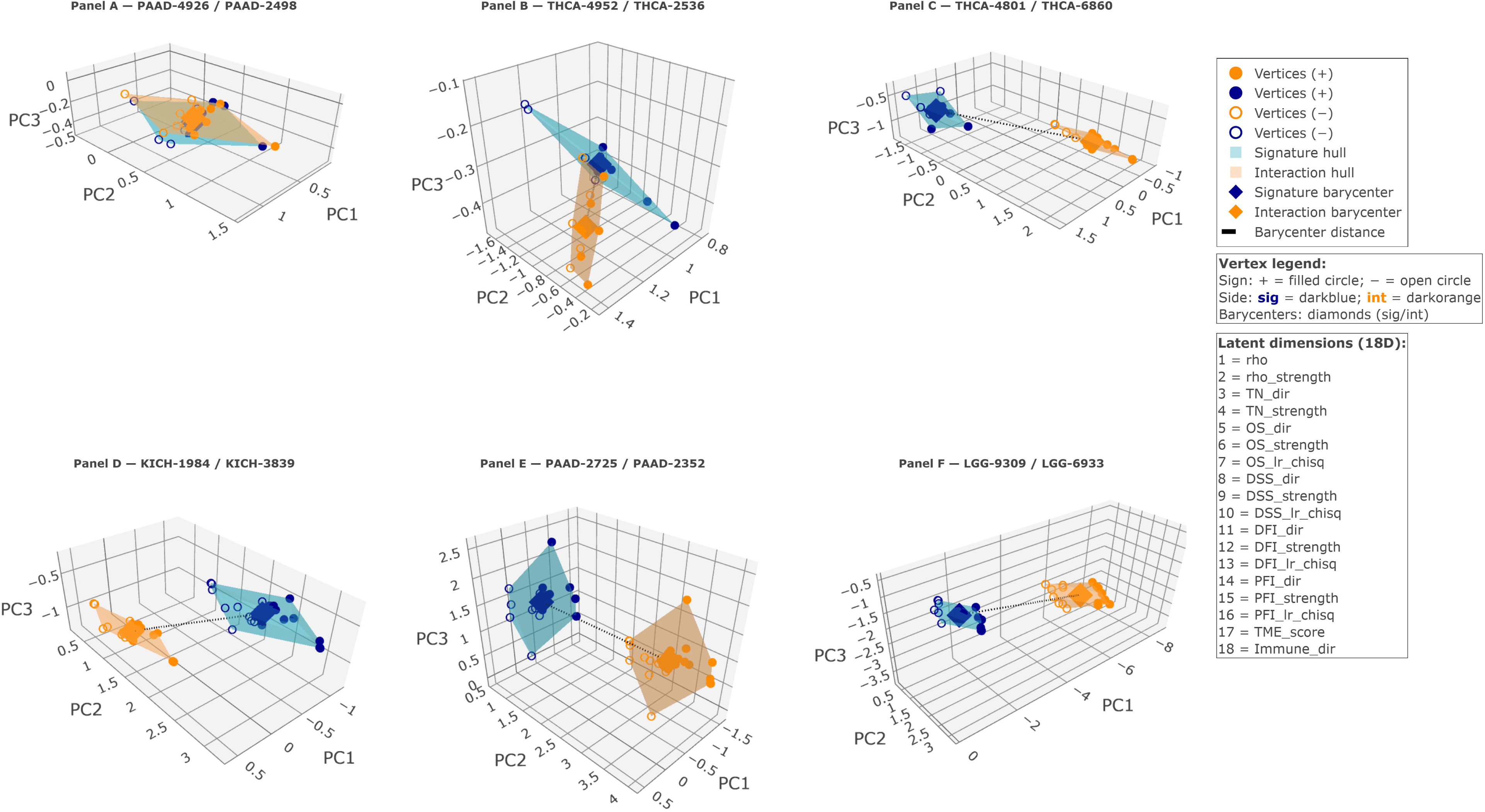
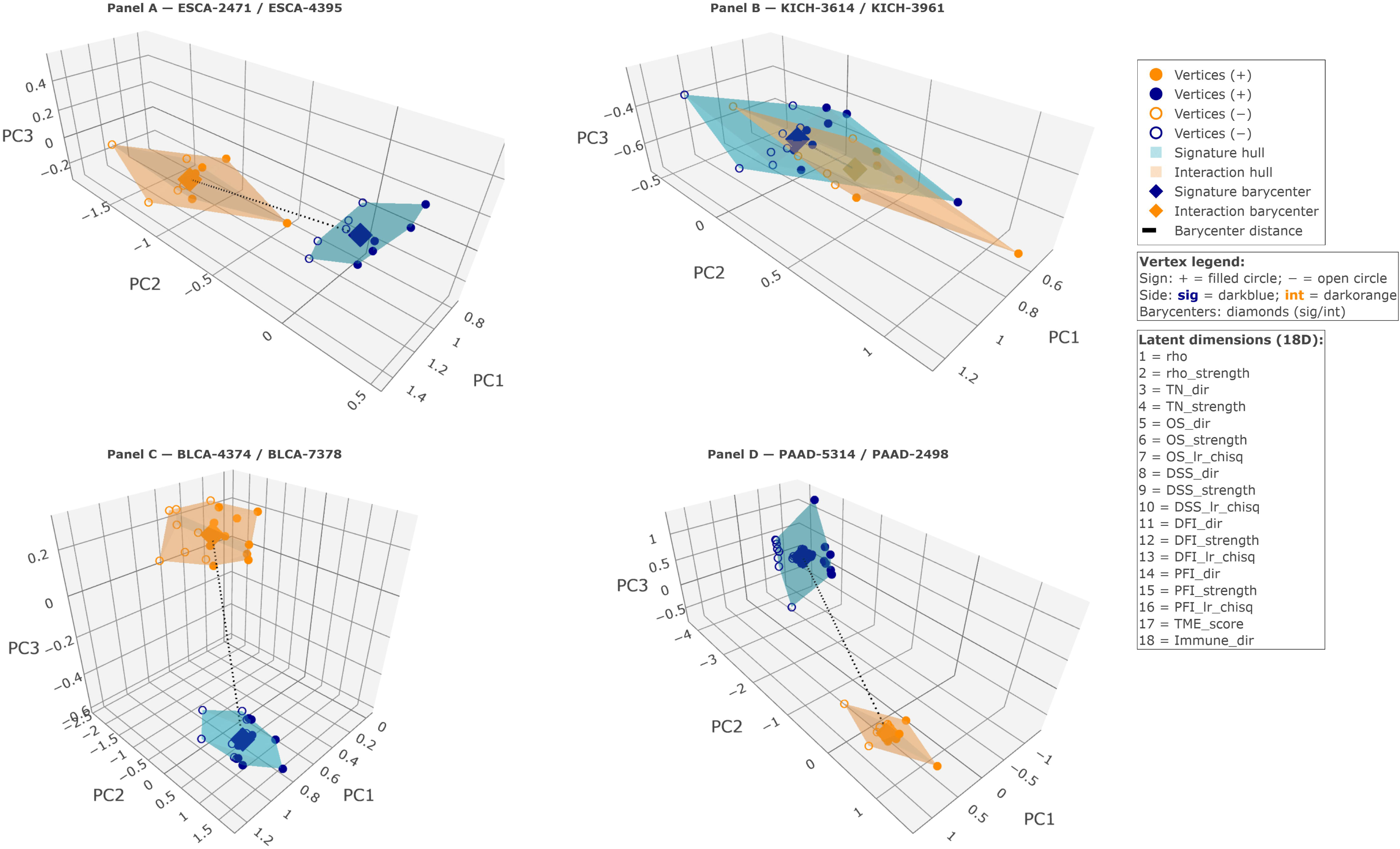

## Notes

https://github.com/BioCancerInformatics/SigPolytope

### Competing Interest Statement

The authors have declared no competing interest.

