## Supplementary Information for "Geometric Multidimensional Representation of Omic Signatures"

**Running title:** Geometry of Multi-Omic Signatures

**Authors:** Higor Almeida Cordeiro Nogueira and Enrique Medina-Acosta

Laboratório de Biotecnologia, Centro de Biociências e Biotecnologia, Universidade Estadual do Norte Fluminense, Brazil

*** Corresponding author:** Enrique Medina-Acosta

**Address:** Laboratório de Biotecnologia, Centro de Biociências e Biotecnologia, Universidade Estadual do Norte Fluminense, Avenida Alberto Lamego 2000, Parque Califórnia, Campos dos Goytacazes, RJ, CEP 28015-602, Brazil**.**

**Authors' e-mails**

**Supplementary Note 1. Nomenclature of signatures and circuitries**

All omic signatures and derived regulatory circuitries in this study follow the standardized nomenclature system introduced in the Oncometabolism GPS framework (1). Each signature is assigned a unique, structured alphanumeric identifier composed of hierarchical tokens encoding tumor type, omic layer, metabolic domain, pathway assignment, phenotypic associations, prognostic behavior, immune context, and regulatory attributes. In the present work, regulatory circuitries are referenced using pairs of these signature identifiers, formatted as Signature A / Signature B, where each component preserves its full nomenclatural encoding. This system ensures lossless traceability between molecular composition, analytical context, and biological interpretation, while maintaining consistency across computational analyses, visualizations, and downstream resources.

**Supplementary Note 2. Definition of Latent Context in the Geometric Framework**

In the present framework, latent context refers to the integrated set of biological, regulatory, and phenotypic constraints that shape the behavior of molecular components but are not directly observable as individual variables. These constraints are not encoded explicitly as predefined annotations or categorical labels. Instead, they emerge from the joint structure of multi-omic associations and are captured through dimensionality reduction into a latent space.

Formally, the latent space constitutes a compressed coordinate system in which each axis summarizes a coherent pattern of covariation across omic layers, phenotypic readouts, and regulatory signals. Individual latent axes do not correspond to single measurable quantities—such as expression levels, mutation states, or pathway memberships—but rather encode integrated contextual effects, including regulatory alignment, phenotypic breadth, tissue specificity, or stability across biological conditions. In this sense, latent context represents the informational environment within which molecular components operate.

When an omic signature is embedded in this latent space, the position of each molecular component reflects not only its individual properties but also its behavior relative to the collective constraints imposed by the system. Components that occupy nearby regions share similar contextual roles, whereas components separated along latent axes diverge in their regulatory, phenotypic, or mechanistic contributions. The latent context therefore defines the background structure against which coordination, divergence, and modularization among components become geometrically apparent.

Importantly, latent context is not inferred a priori from pathway annotations, curated gene sets, or predefined biological categories. It arises from data-driven integration across multiple informational layers and is revealed through the geometry of the embedded signature. The convex hull enclosing the signature captures how molecular components collectively span this latent contextual space, transforming otherwise opaque relational information into an interpretable spatial structure.

**Supplementary Note 3. Construction of the Geometric Representation of Metabolic Regulatory Circuitries**

The complete geometric representation of the metabolic regulatory circuitry dataset was generated by transforming each circuitry into two paired latent objects—one for the signature side and one for the interaction side—and embedding these paired representations into a shared three-dimensional coordinate system that supports direct geometric comparison.

Each circuitry row in the circuitries table was encoded into an 18-dimensional latent vector per side (features_sig and features_int), yielding a dual latent tensor that preserves effect directionality and effect strength across correlation structure, tumor–normal behavior, survival outcomes, and microenvironmental context.

The 18 latent axes were explicitly defined to capture: (i) correlation polarity (rho) and statistical support (rho_strength = −log10(p_adj)); (ii) tumor–normal state as a signed categorical direction (TN_dir, mapped to +1/−1/0 for overexpression, underexpression, or unchanged) together with its evidence strength (TN_strength = −log10(p)); (iii) four survival endpoints (OS, DSS, DFI, and PFI), each decomposed into a Cox direction (dir), a significance-derived strength (strength = −log10(p)), and a log-rank chi-square term (lr_chisq); and (iv) contextual variables comprising a continuous microenvironment score (TME_score) and a signed immune classification (Immune_dir). Robust numerical safeguards were implemented throughout so that missing, non-numeric, or non-positive values were deterministically mapped to zero contribution rather than propagating undefined values into the latent space.

A key design principle enforced by the implementation is that non-significant survival effects do not deform the geometry. Cox-type labels were mapped as Risk → +1, Protective → −1, and NS/NA → 0. Whenever the direction mapped to zero, both the associated strength and chi-square values were explicitly neutralized. As a result, survival endpoints lacking directional support behave as geometrically inactive axes, preventing spurious inflation of polytope extent and ensuring that hull deformation reflects only directionally interpretable effects.

After construction of the paired 18D latent representations, a global principal component analysis (PCA) embedding was computed on the concatenated matrix of all latent vectors from both sides (2N vectors × 18 dimensions). Latent dimensions were centered and scaled using statistics derived from the full combined set, with zero-variance dimensions retained but rendered variance-neutral. This procedure yielded a single shared three-dimensional coordinate system (PC1–PC3) in which both signature-side and interaction-side representations could be compared without introducing side-specific rotations or scaling artifacts.

Each circuitry’s signature and interaction latent vectors were projected into this global PCA space to obtain two barycenters. The Euclidean distance between these barycenters in PCA space was defined as the barycenter distance, which operationalizes a multidimensional discordance magnitude between the two sides of the circuitry. Circuitries were assigned to four barycenter-distance regimes using fixed thresholds in the shared three-dimensional PCA space: high concordance (d_bary_ < 0.5), moderate discordance (0.5 ≤ d_bary_ < 1.5), strong discordance (1.5 ≤ d_bary_ < 2.5), and extreme discordance (d_bary_ ≥ 2.5). These thresholds define the categorical regime labels used throughout the Supplementary tables and analyses and correspond to increasing levels of geometric separation between signature and interaction barycenters in latent space.

To represent not only the barycentric position but also the internal contribution structure of the full 18D vector, each side was expanded into a fixed set of barycenter-centered vertices before convex hull construction. For a latent vector v in R^18, an axis-aligned shell of 2P vertices (P = 18) was generated by perturbing each coordinate v_j by ±step_j, where step_j = max(|v_j|, ε). This yields 36 vertices per side and guarantees non-degenerate steps even when a coordinate is near zero. These vertices were projected into PCA space, and separate convex hulls were computed for the signature and interaction sides. Each circuitry is therefore represented by two co-embedded polytopes plus their barycenters, enabling simultaneous assessment of alignment (via barycenter distance) and geometric complexity (via hull volumes and shape).

For cohort-scale analysis, this geometric construction was evaluated for all circuitries in parallel.

For each circuitry, three primary geometric metrics were extracted: barycenter distance, signature hull volume, and interaction hull volume. From these, the mean hull volume was used as a proxy for latent geometric extent, and a volume ratio (max/min with epsilon guarding) was computed to quantify asymmetry between sides.

Circuitries were assigned to interpretable geometric regimes using deterministic rules. Barycenter distances were discretized into ordered discordance classes ranging from high concordance to extreme discordance using fixed thresholds. Hull-volume complexity strata were defined using data-driven breakpoints based on tertiles of the finite mean volume distribution, with robust fallback behavior when volumes were sparse. These geometric annotations were written back into the full circuitry table with controlled column placement and exported as both RDS and TSV files to preserve linkage with phenotypic and metabolic metadata.

Finally, the geometric classifications served as a scaffold for metabolic overlay analyses. Metabolic pathways and higher-level superfamilies were summarized as distributions across distance-based discordance classes and volume-based complexity and asymmetry classes. For visualization, pathway and metabolism categories were ordered by Shannon entropy of their class distributions, providing an information-theoretic measure of geometric dispersion. This entropy-based ordering replaces arbitrary alphabetic sorting and reveals gradients of metabolic pleiotropy versus specialization across the geometric circuitry landscape.

**Supplementary Note 4. Formal Representation of Omic Signatures as Multidimensional Geometric Objects**

The defining property of an omic signature is its emergence: its biological meaning arises not from any individual feature, but from the organized behavior of all features operating under specific regulatory constraints. Formally, rather than a classical vector s ∈ ℝ^k, an omic signature can be represented as a structured set S = {X₁, X₂, …, Xₙ; Φ}, where each Xᵢ denotes an informational subspace corresponding to a specific molecular, regulatory, or phenotypic modality, and Φ encodes the cross-layer dependencies and conditional relationships governing their coordinated behavior.

Biological instances of Φ include metabolite-driven rewiring of transcriptional programs, microRNA-mediated buffering of metabolic enzymes, context-specific immune modulation of metabolic demand, and stress-induced compensatory pathways characteristic of survival or regulated cell-death processes. In this formulation, the signature is defined by its structural relationships and dynamic organization rather than by its marginal measurements. This structured representation S provides the foundation for encoding omic signatures as point sets within a latent space.

The geometric properties of an omic signature further distinguish it from traditional vectorial representations. Each informational subspace may occupy a distinct metric space and possess its own dimensionality, and their integration yields a structure more appropriately conceived as a multi-layered manifold rather than a single point in Euclidean space. In this view, each omic layer contributes a fiber, their interactions define the connections among fibers, and the emergent phenotype corresponds to the curvature of the manifold within specific regions of this space.

Two signatures that appear similar when reduced to gene lists or composite scores may nevertheless exhibit profoundly different manifold geometries, reflecting distinct patterns of regulatory organization, inter-layer coupling, or contextual constraint.

Within this multidimensional formulation, a faithful mathematical representation must capture both the magnitude and the structural organization of an omic signature across its interacting dimensions. Such an entity can be conceptualized as a convex polytope embedded within an n-dimensional latent space, where each axis summarizes a biologically meaningful latent variable such as molecular effect, regulatory breadth, phenotypic dispersion, or contextual stability.

This perspective motivates the use of geometric constructs—such as convex hulls—to preserve the multidimensional structure of biological variation. Convex-hull–based polyhedral representations have demonstrated utility in multidimensional biomedical phenotyping, including human systems stratification (2). However, most prior implementations are limited to three-dimensional instantiations of a broader conceptual framework.

In the present formulation, omic signatures are treated as intrinsically n-dimensional informational polytopes whose convex hulls capture their structural identity and delineate the boundaries of their informational coherence within latent biological space. The convex polytope—or its observable three-dimensional projection as a convex hull—serves as a minimal, topology-preserving envelope encompassing all constituent components of an omic signature. Intuitively, the convex hull corresponds to the smallest surface enclosing all points, while formally representing the smallest convex set containing them. This representation preserves the internal organization of the signature while enabling quantitative geometric comparison across signatures, contexts, and biological systems.

### **Supplementary Note 5. Formal Geometric Representation of Omic Signatures**

Formally, an omic signature can be represented as a point set S = {x₁, x₂, …, xₙ} ⊂ ℝⁿ, where each xᵢ corresponds to a molecular component mapped to latent coordinates summarizing its multi-omic associations. Endowing ℝⁿ with a metric structure allows distances, angles, and topological relationships among components to be computed, enabling the signature to be treated as a quantifiable geometric object rather than a list of molecules.

Within this space, the convex hull Conv(S)—defined as the smallest convex set containing all elements of S—captures the structural boundaries of the signature and constitutes the minimal envelope that preserves its multidimensional organization. Although signatures reside intrinsically in ℝⁿ, a projection P: ℝⁿ → ℝ³ is employed to obtain a three-dimensional representation suitable for visualization, while retaining the salient geometric properties required for analysis.

The geometric properties of this representation provide intrinsic measurements of signature structure. Hull volume serves as a proxy for multidimensional amplitude, principal axes derived from the covariance structure of S capture dominant sources of mechanistic variation, and anisotropy quantifies the degree to which the signature is driven by a single coherent process versus multiple partially orthogonal subprograms. Additional descriptors, including centroid location, diameter, surface morphology, and angular divergence between hulls, support systematic comparison and classification.

These formal definitions provide the mathematical foundation for the geometric framework employed throughout the study and underlie all quantitative analyses reported in the main text.

**Supplementary figure legends**

**Figure S1. Representative convex-hull geometries across observed latent complexity × symmetry states** ([FS1_HTML](https://biocancerinformatics.github.io/SigPolytope/figures/html/Figure_S1.html))**.**

Convex hulls were constructed from barycenter-centered perturbations along all 18 latent axes to resolve local multidimensional structure around each regulatory circuitry. Hull volume provides a proxy for latent complexity, stratifying circuitries into low-dimensional (flat), intermediate-complexity, and high-complexity multidimensional geometries. Symmetry was assessed using the volume-implication regime encoded in vol_implication. Panels are arranged as a 3 × 2 grid (rows: latent complexity tier; columns: symmetric vs signature-dominant asymmetry). Panels A–B show low-dimensional geometries (symmetric and signature-dominant, respectively), Panels C–D show intermediate-complexity geometries (symmetric and signature-dominant), and Panels E–F show high-complexity multidimensional geometries (symmetric and signature-dominant). Asymmetry magnitude (moderately vs strongly asymmetric) is pooled into a single signature-dominant category for visualization clarity; quantitative distributions of all volume-implication regimes are reported in Table 1B. For each panel, a population-level representative circuitry was selected as the median-central geometry within its class based on (sig_hull_vol, int_hull_vol, vol_ratio). No interaction-dominant geometries were observed in Dataset S1. Panel–circuitry mapping and geometric descriptors for Figure S1 are provided in Supplementary Table S1.

**Figure S2. Population-level exemplars of geometric regulatory circuitries illustrating the full spectrum of structural behaviors observed across the atlas** ([FS2_HTML](https://biocancerinformatics.github.io/SigPolytope/figures/html/Figure_S2.html)).

Figure S2 presents representative, empirically observed circuitries selected from the full atlas to illustrate the range of geometric phenotypes detected at the population level. Circuitries are represented as paired convex polytopes corresponding to signature and interaction profiles, embedded in a shared 18-dimensional latent space and projected onto a common three-dimensional PCA coordinate system. Geometric discordance is quantified by the Euclidean distance between polytope barycenters, while internal structural organization is captured by convex-hull volumes and their ratio, reflecting relative dimensional complexity and symmetry. Biological concordance is evaluated across phenotypic, immune, and survival (hazard ratio–based) dimensions. Panel A. Strongly discordant, low-dimensional asymmetric circuitry (ESCA). A strongly discordant circuitry linking hsa-miR-17-5p (ESCA-2471) and RRM2 (ESCA-4395) is shown (barycenter distance = 1.53). The geometry is highly asymmetric, with a compact, flattened signature polytope (hull volume = 3.8 × 10⁻³) contrasted against an expanded interaction polytope (hull volume = 5.27 × 10⁻²; volume ratio = 13.85). The signature (miRNA expression, stemness) is overexpressed, anti-tumoral, and cold immune, with no meaningful survival association, whereas the interaction (DNA methylation, stemness) exhibits inverse internal correlation, a pro-tumoral microenvironment, and a meaningful protective survival implication. Immune classification converges, but phenotypic interpretations diverge (Immune: Convergent; Phenotype: Divergent). This circuitry maps to nucleotide metabolism, specifically purine metabolism. Panel B. Highly concordant, symmetric low-dimensional circuitry (KICH). A highly concordant circuitry connecting hsa-miR-98-5p (KICH-3614) and QDPR (KICH-3961) shows close barycenter alignment (barycenter distance = 0.30) and near-equal polytope volumes (signature = 5.91 × 10⁻²; interaction = 5.11 × 10⁻²; volume ratio = 1.16). Both sides are stemness-associated, exhibit coherent negative internal correlation, share a cold immune state, and display meaningful risky survival associations, yielding overall concordance across phenotypic and immune dimensions. This circuitry maps to metabolism of cofactors and vitamins, specifically folate biosynthesis. Panel C. Extremely discordant circuitry with intermediate complexity (BLCA). An extremely discordant circuitry linking hsa-miR-193b-3p (BLCA-4374) and MPST (BLCA-7378) exhibits a large barycenter separation (barycenter distance = 2.73). Both polytopes display intermediate internal structure (signature hull volume = 5.23 × 10⁻²; interaction = 1.16 × 10⁻¹; volume ratio = 2.22), indicating moderate asymmetry without dimensional collapse. Although both sides are stemness-associated and overexpressed, survival implications diverge (risky vs. protective), and immune context is discordant (hot vs. cold), resulting in Immune: Divergent; Phenotype: Convergent; Cox: Divergent. This circuitry maps to energy metabolism, specifically sulfur metabolism. Panel D. Extremely discordant circuitry dominated by complexity asymmetry (PAAD). An extreme discordance exemplar connecting hsa-miR-376c-5p (PAAD-5314) and GGCX (PAAD-2498) shows the largest barycenter separation among the panels (barycenter distance = 4.53). Discordance is driven by pronounced differences in internal complexity, with a highly multidimensional signature polytope (hull volume = 0.82) opposed to a compact interaction polytope (hull volume = 5.28 × 10⁻²; volume ratio = 15.47). Despite shared cold immune context, phenotypic and survival interpretations diverge, with protective vs. risky survival associations (Immune: Convergent; Phenotype: Divergent; Cox: Divergent). This circuitry maps to metabolism of cofactors and vitamins, specifically ubiquinone and other terpenoid-quinone biosynthesis.

**Figure S3. Entropy-ordered distribution of geometric distance-implication classes across Final Concordance Categories.** The figure displays the proportional distribution of circuitries across the four geometric distance-implication regimes (high concordance, moderate discordance, strong discordance, and extreme discordance) for each *Final_concordance_summary* category. Categories are ordered from highest to lowest Shannon entropy computed over the distribution of geometric regimes, such that classes exhibiting the greatest geometric heterogeneity appear first. Entropy quantifies the dispersion of circuitries across geometric states and serves as a measure of structural diversity within each concordance class. Higher-entropy classes, such as *Only Convergent*, populate a wide range of geometric configurations, whereas lower-entropy classes, exemplified by *Only Divergent*, concentrate in fewer geometric regimes. This ordering reveals that biologically divergent circuitry states tend to arise from more structurally constrained geometric configurations, while biologically concordant states emerge from a broader and more permissive region of the geometric space. Together, the entropy profile and proportional distributions highlight distinct geometric architectures underlying convergence and divergence in immune, phenotypic, Cox, and survival concordance dimensions.

**Supplementary Table S1 Figure S1 panel key.** Mapping between Figure S1 panels (A–F) and the selected representative circuitries, including hull volumes, volume ratio, and the corresponding volume-implication regime.

| **Panel** | **Panel_description** | **Circuitries_id** | **Geometry_class** | **Complexity_tier** | **Symmetry_state** | **sig_hull_vol** | **int_hull_vol** | **vol_ratio** | **dist_to_median** | **vol_implication** |
| --- | --- | --- | --- | --- | --- | --- | --- | --- | --- | --- |
| A | Low — Symmetric | PAAD-4926 / PAAD-2498 | Low-dimensional, symmetric geometry | Low | Symmetric | 6.079e-02 | 5.278e-02 | 1.152e+00 | 4.183e-02 | low_dimensional_flat ; symmetric_sig_int |
| B | Low — Signature-dominant | THCA-4952 / THCA-2536 | Low-dimensional, signature-dominant asymmetric geometry | Low | Sig_dominant | 1.671e-02 | 9.317e-04 | 1.794e+01 | 8.925e-01 | low_dimensional_flat ; strongly_asymmetric_sig_int |
| C | Intermediate — Symmetric | THCA-4801 / THCA-6860 | Intermediate-complexity, symmetric geometry | Intermediate | Symmetric | 1.283e-01 | 1.082e-01 | 1.186e+00 | 4.984e-02 | intermediate_complexity ; symmetric_sig_int |
| D | Intermediate — Signature-dominant | KICH-1984 / KICH-3839 | Intermediate-complexity, signature-dominant asymmetric geometry | Intermediate | Sig_dominant | 1.876e-01 | 8.003e-02 | 2.344e+00 | 2.746e-01 | intermediate_complexity ; moderately_asymmetric_sig_int |
| E | High — Symmetric | PAAD-2725 / PAAD-2352 | High-complexity multidimensional, symmetric geometry | High | Symmetric | 6.974e-01 | 8.081e-01 | 1.159e+00 | 3.746e-02 | high_complexity_multidimensional ; symmetric_sig_int |
| F | High — Signature-dominant | LGG-9309 / LGG-6933 | High-complexity multidimensional, signature-dominant asymmetric geometry | High | Sig_dominant | 2.372e-01 | 1.445e+00 | 6.092e+00 | 5.984e-01 | high_complexity_multidimensional ; strongly_asymmetric_sig_int |

**Repository links to interactive HTML figures**

The links below point to interactive HTML figures hosted on GitHub Pages that were used during analysis and exploratory validation of geometric structures. These HTML figures are provided for transparency and interactive inspection only. They are not publication-ready figures and should not be considered authoritative renderings.

All figures appearing in the main manuscript and Supplementary Information were regenerated directly from the underlying data using deterministic, non-interactive plotting pipelines and exported as vector graphics prior to conversion to high-resolution TIFF/PDF formats. As a result, the final publication figures may differ in layout, styling, or rendering details from the interactive HTML versions.

| **Figures** | **GitHub link** |
| --- | --- |
| Figure 2_HTML | <https://biocancerinformatics.github.io/SigPolytope/figures/html/Figure_2.html> |
| Figure 3_HTML | <https://biocancerinformatics.github.io/SigPolytope/figures/html/Figure_3.html> |
| Figure 4_HTML | <https://biocancerinformatics.github.io/SigPolytope/figures/html/Figure_4.html> |
| Figure S1_HTML | <https://biocancerinformatics.github.io/SigPolytope/figures/html/Figure_S1.html> |
| Figure S2_HTML | <https://biocancerinformatics.github.io/SigPolytope/figures/html/Figure_S2.html> |
